# 5’ modifications to CRISPR Cas9 gRNA can change the dynamics and size of R-loops and inhibit DNA cleavage

**DOI:** 10.1101/2020.04.09.033399

**Authors:** Grace Mullally, Kara van Aelst, Mohsin M. Naqvi, Fiona M. Diffin, Tautvydas Karvelis, Giedrius Gasiunas, Virginijus Šikšnys, Mark D. Szczelkun

**Affiliations:** DNA-Protein Interactions Unit, School of Biochemistry, University of Bristol, BS8 1TD, UK; BrisSynBio, Life Sciences Building, Tyndall Avenue, University of Bristol, Bristol, BS8 1TQ, UK; Institute of Biotechnology, Vilnius University, Vilnius, Lithuania; CasZyme, Vilnius, Lithuania

## Abstract

A key aim in exploiting CRISPR-Cas is the engineering of gRNA to introduce additional functionalities, ranging from small nucleotide changes that increase efficiency of on-target binding to the inclusion of large functional RNA aptamers and ribonucleoproteins (RNPs. Interactions between gRNA and Cas9 are crucial for RNP complex assembly but several distinct regions of the gRNA are amenable to modification. Using a library of modified gRNAs, we used *in vitro* ensemble and single-molecule assays to assess the impact of RNA structural alterations on RNP complex formation, R-loop dynamics, and endonuclease activity. Our results indicate that R-loop formation and DNA cleavage activity are essentially unaffected by gRNA modifications of the Upper Stem, first Hairpin and 3’ end. In contrast, 5’ additions of only two or three nucleotides reduced R-loop formation and cleavage activity of the RuvC domain relative to a single nucleotide addition. Such gRNA modifications are a common by-product of *in vitro* transcribed gRNA. We also observed that addition of a 20 nt RNA hairpin to the 5’ end supported formation of a stable ~9 bp R-loop that could not activate DNA cleavage. These observations will assist in successful gRNA design.

## INTRODUCTION

The microbial CRISPR-Cas systems, and in particular type II-A Cas9, have found widespread utility as tools for site-specific DNA and RNA recognition (1,2). They can be readily programmed by changing the spacer sequence of a crRNA that recognises a DNA target protospacer by forming an R-loop, binding to the targeted strand (TS) through Watson-Crick base pairing (3). DNA recognition also requires binding of a protospacer adjacent motif (PAM), e.g. *Streptococcus pyogenes (Sp)* Cas9 binds 5’-NGG-3’ while *Streptococcus thermophilus* DGCC7710 CRISPR3 *(St3)* Cas9 binds 5’-NGGNG-3’ (4,5). Cas9 evolved to use a trans-activating RNA (tracrRNA) that base pairs via an anti-repeat sequence to the repeat sequence of individual crRNAs (4,6). The single gRNA more commonly used in tool applications is a fusion of the tracrRNA and crRNA through a short RNA loop between the repeat-anti-repeat sequences in the Upper Stem (Figure 1) (4). An important area of research has been the manipulation of the gRNA sequence and structure, through introduction of extra or modified nucleotides, and/or the addition of folded RNA aptamers (7,8). Using a combination of ensemble and single-molecule biochemical assays, we examined how gRNA modification affects ribonucleoprotein (RNP) assembly, R-loop stability and DNA cleavage. Our results indicate that *Sp*Cas9 R-loop formation and DNA cleavage activity is acutely sensitive to changes to the 5’ end of the RNA, even when the addition is only two nucleotides.

**Figure 1.**
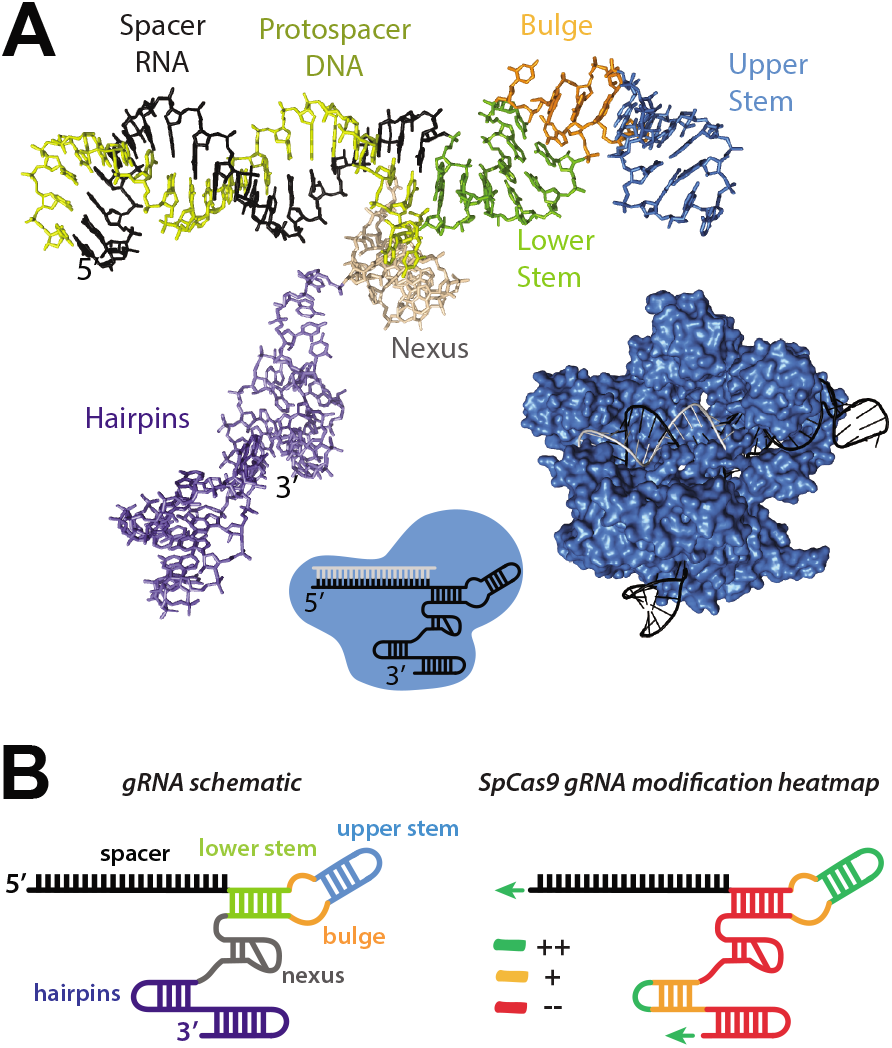
*Streptococcus pyogenes* Cas9 gRNA structure. (**A**) The structure of *Sp*Cas9 (surface render in blue) in complex with gRNA (black) and target DNA (grey) from PDB 4OO8 (9). The gRNA and protospacer DNA structures are shown separately with structural features labelled: spacer (black); Lower Stem (green); Bulge (orange); Upper Stem (blue); Nexus (beige); Hairpins (purple). (**B**) *(left)* gRNA schematic as used in other figures with colouring consistent with panel B. *(right)* gRNA heatmap, based on Briner et al (10), showing gRNA regions which tolerate (green), partially tolerate (orange) and do not tolerate (red) modification.

The highly-folded Lower and Upper Stems, Nexus and Hairpin regions of the gRNA make extensive contacts with positively-charged surfaces of Cas9 (Figure 1A): Cas9 folds around the RNA/DNA duplex and the Lower Stem-Bulge-Upper Stem region using the recognition (REC) and nuclease (NUC) lobes; much of the hairpin regions of the gRNA contact a positively-charged region on the Cas9 outer surface; the gRNA provides most of the interactions between the REC and NUC lobes which themselves do not form extensive protein-protein interactions. Removing structures with extensive Cas9-contacts, such as the Bulge and Nexus, abolished Cas9 function while mutations were tolerated in other regions of the gRNA, particularly those not directly contacted by Cas9 such as the Upper Stem (9). An extensive study by Briner *et al* confirmed the importance of the Bulge and Nexus and also highlighted the possibility of modifying the 5’ and 3’ termini, Upper Stem and first Hairpin (10). These studies can be summarised as a theorised ‘heatmap’ of modification tolerance, Figure 1B.

Modification of the 5’ end of gRNAs is a familiar by-product of the everyday use of Cas9 in gene editing. T7 bacteriophage RNA polymerase is commonly used for *in vitro* transcription (IVT) of gRNAs. The T7 promoter has been minimised for *in vitro* use; whereas naturally occurring initiation sequences have up to 3 guanines in the +1, +2 and +3 positions (11), the minimal requirement for initiation is one guanine in the +1 position resulting in a single guanine at the 5’ end of the spacer sequence. Nonetheless, two or more guanines are often recommended to ensure higher yields of transcribed gRNA. Commonly used mammalian promoters for cellular gRNA transcription have similar initiation requirements; gRNAs expressed from the U6 promoter necessitates a G or A in the (+1) position for high expression. By careful choice of protospacer target sequence, these nucleotides can be incorporated into the spacer and thus R-loop DNA-RNA hybrid. However, in many cases this is not feasible, and the additional nucleotides become an unpaired overhang.

Unpaired nucleotides at the 5’ end of the spacer sequence can influence *SpCas9* activity in a protein and target site-dependent manner (12,13). For WT Cas9, one or two additional 5’ guanines lead to increased specificity due to reduced unwinding promiscuity. However, two 5’ guanines caused a decrease in on-target activity. For the enhanced SniperCas9, one unpaired 5’ guanine had a positive effect in increasing the sensitivity to mismatches but two unpaired 5’ guanines lowered specificity. Other engineered Cas9s showed either reduced on-target activity (eCas9 and HypaCas9) or increased promiscuity (Cas9-HF1). These pleiotropic effects are most likely due to the observed interactions of the docked RuvC domain with the 5’ end of the RNA (9,13). Unpaired 5’ nucleotides could produce distortion of the DNA:RNA hybrid in this region that may influence R-loop formation, and possibly DNA cleavage.

There are also numerous examples of gRNA that have been had additional functional RNA structures appended [(reviewed in (7)], including ribozymes at the 5’ end (12,14), modification of the upper stem with RNA aptamer binding effectors to colocalise fluorescent proteins (15), and 3’-fusion of viral RNA scaffolds to recruit a variety of transcriptional activators, suppressors, or protein modifiers for reprogramming gene expression (16). It is perhaps surprising given the sensitivity to even a single unpaired nucleotide that larger RNA structures can successfully accommodated at the 5’ end of Cas9 gRNAs [e.g. (17)]. Although using gRNAs as RNA-mediated scaffolds has been broadly shown to allow functionality in the downstream applications, generally in cells, little work has been done to understand the impacts of adding RNA scaffolds to gRNA on CRISPR RNP complex assembly, R-loop formation/dissociation, and downstream activation of nuclease function (where used). Knowledge of how gRNA-fusions affect all Cas9 activities will contribute to the rationale design of these structures.

We previously developed a single-molecule magnetic tweezers (MT) assay to monitor R-loop formation by *St3*Cas9 with its natural crRNA:tracrRNA (18). We used this assay here with both *St3*Cas9 and *Sp*Cas9 to explore the effects of gRNA modifications on R-loop formation. We additionally used a FRET-based assay to measure formation of the RNP with each gRNA (19), and the effect on the rate of dsDNA cleavage using a supercoiled plasmid substrate where we can monitor formation of nicked intermediate and cleavage product without R-loop formation being rate-limiting (20). We show that 2 or 3 unpaired 5’ guanine or uracil nucleotides result in inhibition of cleavage of the non-targeted strand by the RuvC nuclease and in some cases alterations in the R-loop formation dynamics. Our results are consistent with complex and variable effects that are influenced by RNA and/or local DNA sequence.

We additionally tested the effects of a 20 nt structured RNA hairpin [the putative mitochondrial targeting RNA aptamer, RP – (21)] when added to the 5’ end, Stem Loop, first Hairpin or 3’ end of the gRNA of *Sp*Cas9. Except for the 5’ modification, the RP hairpin had only moderate effects on R-loop formation dynamics and no measurable effects on the DNA cleavage kinetics. In contrast, the addition of the RP structure to the 5’ end resulted in formation of a stable half-sized R-loop that did not activate DNA cleavage. We propose that the structured RNA cannot be threaded into a full-length R-loop but can still produce a stable structure but where the RuvC/HNH domains are not docked in an active conformation. These observations highlight that modifications to other parts of the gRNA, most prudently the 3’ end, will be more likely to be successful if DNA cleavage is required but that 5’ modification could be used where just DNA binding is required.

## MATERIALS and METHODS

### gRNAs

IVT dsDNA templates were amplified by PCR from the relevant plasmid (see Supplementary Table 2 for sequences): pD1301i-SP1 was generated by cloning the 1025-1044 pSP1 spacer sequence into pD1301i (Atum Bio); the pEX-A2 family of plasmids were supplied by Eurofins; pCRISPR3 was from ref (22); pX330 was a gift from Feng Zhang (Addgene plasmid # 42230; http://n2t.net/addgene:42230; RRID:Addgene_42230) (23). Primers were designed for IVT that amplified the RNA spacer and structural component and introduced the T7 promoter upstream of the gRNA sequence. IVT was performed with HiScribe T7 High Yield RNA synthesis kit (New England BioLabs) as per the manufacturer’s instructions. RNAs were purified by either phenol-chloroform extraction or with RNA Clean and Concentrator Columns (Zymo Research) and eluted in DEPC treated H2O. Synthetic crRNAs and gRNAs were supplied by Metabion and IDT, respectively (Supplementary Figure S1). gRNA concentrations were calculated by taking A_260_ with a DeNovix DS-11 spectrophotometer.

### Cas9 Protein

Wild type *St*3Cas9 was expressed and purified as published previously (6). *St*3Cas9(D31A), *St*3Cas9(N891A) and *St*3dCas9 variants were engineered from pBAD-Cas9 plasmid (6) using a Phusion Site-Directed Mutagenesis Kit (Thermo Fisher Scientific), and expressed and purified as for WT *St3Cas9. Sp*Cas9 was either supplied by New England Biolabs or the *Sp*Cas9 encoding gene was cloned into pBAD (6) from pMJ806 (4) and expressed/purified as for WT St3Cas9.

A plasmid to express *Sp*Cas9_hinge_ (19) was produced by cloning five synthetic gene fragments in pE-SUMO (Life Sensors, PA) using NEBuilder HiFi DNA Assembly (New England Biolabs). For purification, *E. coli* BL21 Rosetta 2 (DE3) cells were transformed with pSUMOCas9_hinge_ and grown overnight on LB with 50 μg/ml ampicillin. 500 ml LB was inoculated with a single colony and cells were grown at 37 °C in a 2.5 L flask at 250 rpm for 16 hours. Once grown, bacteria were pelleted by centrifugation for 20 mins at 4,000 x g, 4 °C, and supernatant discarded. Pellets could be frozen in liquid nitrogen and stored at −20 °C at this point. The bacterial pellet was resuspended in 25 ml sonication buffer [50 mM Tris, pH 8.0, 500 mM NaCl, 1 mM β-mercaptoethanol, 5 mM MgCl_2_, 0.5 mM EDTA, Roche cOmplete ULTRA protease inhibitor] and lysed by sonication 2 x 2 minutes (10 seconds on/off) at 75% maximum power. Total lysate was cleared by ultracentrifugation for 40 minutes at 37,000 x g, 4 °C and supernatant was dialysed at 4 °C against 1L HisWash1 [50 mM Tris, pH 9.0, 500 mM NaCl, 1 mM β-mercaptoethanol, 30 mM imidazole] in Snakeskin 10,000 MWCO Dialysis tubing (Thermo Scientific). Sample was filtered through a 0.45 μm nitrocellulose filter and loaded onto a 5 ml HisWash1-equilibrated HisTrap HP column (GE Healthcare). 2 ml fractions were collected through a linear gradient over 100ml up to 100% HisElution [50 mM Tris, pH 9.0, 500 mM NaCl, 1 mM β-mercaptoethanol, 500 mM imidazole]. Cas9_hinge_ eluted in the range of 120 – 170 mM imidazole. Pooled fractions were dialysed at 4 °C against HisWash2 [50 mM Tris, pH 9.0, 200 mM NaCl, 1 mM β-mercaptoethanol] and total protein concentration estimated using a DeNovix DS-11 spectrophotometer. The His tag-SUMO was cleaved overnight with SUMO protease (1 mg SUMO protease was added per mg Cas9 protein) in Snakeskin dialysis tubing, dialysed against 2 L HisWash2 at 4 °C. After 15 hours, an appropriate amount of NaCl and imidazole were added to give a final concentration of 30 mM imidazole, and the cleaved sample was loaded onto a HisWash2 equilibrated 5 ml HisTrap HP column (GE Healthcare). The flow-through was collected as 2 ml fractions, and bound tag and SUMO protease eluted with HisElution. Cas9-containing fractions were concentrated and exchanged into GF buffer [20 mM Tris, pH 7.5, 200 mM KCl, 1 mM TCEP, 10% (v/v) glycerol] with Amicon Ultra-15 50 kDa cut-off centrifugal filter units (Millipore), and frozen with liquid nitrogen and stored at – 80°C.

*Sp*Cas9_hinge_ was either single labelled with Cy3 only or double labelled with Cy3 and Cy5 using maleimide monoreactive dyes (Amersham, GE Healthcare) (19). 10 μM *Sp*Cas9_hinge_ and 25 μl dye in GF buffer were incubated for 2 hours at room temperature, and overnight at 4 °C. The reaction was quenched with 10 mM DTT and free dye was separated by size exclusion chromatography on a HiLoad Superdex 200 16/60 column (GE Healthcare). Fractions contained labelled protein were pooled and concentrated with Amicon Ultra-15 50 kDa cutoff centrifugal filter units (Millipore), frozen with liquid nitrogen and stored at −80°C. To assess labelling efficiency, proteins were diluted and scanned in a quartz cuvette with a Cary 60 UV-Vis spectrophotometer (Agilent). Extinction coefficients for Cas9, Cy3 and Cy5 at 280 nm (protein), 552 nm (Cy3) and 650 nm (Cy5) were used to calculate the protein concentration and labelling efficiency.

### Ribonucleoprotein assembly

Unless otherwise stated, for RNP complex assembly, equal concentrations of Cas9 and gRNA were mixed in SB buffer [10 mM Tris-Cl, pH 7.5, 100 mM NaCl, 1 mM EDTA, 0.1 mM DTT, 5 μg/ml BSA] for binding assays, or RB buffer [10 mM Tris-Cl, pH 7.5, 100 mM NaCl, 10 mM MgCl_2_, 0.1 mM DTT, 5 μg/ml BSA] for cleavage assays, and incubated at 37 °C for 1 hour, and then at 4°C.

### FRET Measurements of gRNA Loading

100 nM gRNA was assembled with 50 nM *Sp*Cas9_hinge_ labelled with Cy3 or Cy3/Cy5 on ice. Following a 10 min incubation at room temperature, measurements were collected using a FluoroLog Spectrophotometer (Horiba) in a 130 μl Quartz cuvette (Hellma); 5 mm slit widths, 1 s integration time. For each RNA, the sample was excited at 530 nm and 630 nm and spectra collected from 550 – 800 nm and 650 – 800 nm respectively. (Ratio)_A_, calculated as explained in Supplementary Figure 4 as a proxy measure for FRET (19).

### Single Molecule Magnetic Tweezers Assay

The magnetic tweezers assays used a commercial PicoTwist microscope (Fleurieux sur L’Arbresle, France) equipped with a 60 Hz Jai CV-A10 GE camera (24). DNA molecules were tethered to 1 μm MyOne paramagnetic beads (Invitrogen) and the glass coverslip of the flow cell as previously described (18). Topologically-constrained DNA [pSP1, (22)] were identified from rotation curves at 0.3 pN and the rotational zero reference (Rot0) set. R-loop dynamics were measured in Buffer SB at 25 C using magnet rotations of 10 turns s^-1^. The R-loop size in turns was estimated by comparing the slope of the rotation curves at 1 turns s^-1^ when an R-loop was trapped in negative torque with the equivalent slopes in the absence of enzyme as described in van Aelst et al (20). Each trace in a reference set of curves collected in the absence of enzyme (a total of 20) was compared to each rotation curve when an R-loop was trapped in the presence of enzyme (a total of 22). Torque values were calculated using software described in ref (18).

### DNA cleavage Assays

For each cleavage reaction, 3 nM plasmid substrate pSP1 in RB buffer was pre-heated at 20 °C for 5 minutes. The reaction was started by addition of 50 nM assembled Cas9 RNP and incubated for the time period specified. The reaction was quenched by adding 0.5 volumes of STEB [0.1 M Tris (pH 7.5), 0.2 M EDTA, 40% (w/v) sucrose, 0.4 mg/ml bromophenol blue] and incubating at 80 °C for 5 minutes. Samples were separated by agarose gel electrophoresis on a 1.5 % (w/v) agarose gel in 1X TAE [40 mM Tris-acetate, 1 mM EDTA, 10 μg/ml ethidium bromide] at 2 V/cm overnight (16 hours) and visualised by UV irradiation. DNA bands containing supercoiled, linear or open circle DNA were excised and placed into scintillation vials. 0.5 ml sodium perchlorate was added to each gel slice, and tubes were incubated at 67°C for 2 hours to melt the agarose. The vials were cooled to room temperature and 10 ml Hionic-Fluor Scintillation Cocktail (Perkin Elmer) added to each vial and shaken thoroughly. Each vial was counted in a Tri-Carb Trio 3100TR Liquid Scintillation Counter for 10 minutes. Where indicated, the cleavage data was fitted to the models in Supplementary Figure 6 using numerical integration in Berkeley Madonna (www.berkeleymadonna.com).

## RESULTS

### Similar R-loop formation dynamics of wild type and nuclease mutants of *Streptococcus thermophilus* and *Streptococcus pyogenes* Cas9 using crRNA:tracrRNA

In our previous MT study of R-loop formation, we utilised the type II-A *St3*Cas9 and its natural crRNA:tracrRNA (18). Here we repeated the MT assay to compare R-loop formation by *St3*Cas9 with *Sp*Cas9 using their corresponding crRNA:tracrRNA (Supplementary Figures 1 and 2, Supplementary Tables 1 and 2). These are highly related Cas9s, which can even exchange their crRNA:tracrRNA *in vitro,* so we anticipated similar dynamics. crRNAs were synthesised using phosphoramadite chemistry and thus the 5’ end was paired in the R-loop (Supplementary Figure 2A). tracrRNAs were synthesised by *in vitro* transcription (IVT) using T7 RNA polymerase, resulting in 3 unpaired 5’ guanines (Supplementary Figure 1 and Supplementary Figure 2A). These are sited at the end of the Upper Stem where they are free of Cas9 interaction. We also measured the R-loop formation using the nuclease dead (D31A, N891A) and nicking variants (RuvC mutant D31A and HNH mutant N891A) of *St3*Cas9 (5). Some restriction enzymes mutated within their active sites have tighter DNA binding activity than WT enzymes.

Linear DNA derived from pSP1 with a shared protospacer and overlapping *St3* (5’-NGGNG-3’) and *Spy* (5’-NGG-3’) PAMs (Supplementary Figure 2A), was tethered between a glass coverslip and a paramagnetic bead (Supplementary Figure 2B). The bead position above the glass surface was monitored at 60 Hz using video microscopy. Using a pair of permanent magnets, the DNA was first negatively supercoiled, favouring DNA unwinding and R-loop formation. This caused the bead height to lower. The binding and unwinding of the DNA by Cas9 caused a reduction in DNA twist which was compensated by an increase in twist and writhe elsewhere in the DNA. This reduced the negative supercoils and the bead height increased. The magnets were then rotated rapidly to form positive supercoils, conditions favouring DNA rewinding and R-loop dissociation. The increase in DNA twist that accompanied R-loop dissociation was compensated by a reduction in DNA twist/writhe. This reduced the positive supercoils and the bead height increased. The reaction buffer contained EDTA to prevent DNA cleavage so that this cycle could be repeated *ad infinitum*. The force was varied to explore the effect of torque, as required.

Each of the enzymes showed the characteristic changes in bead height corresponding to formation and dissociation of a 20 bp R-loop (Supplementary Figure 2C). R-loop formation was irreversible over several hours and could only be dissociated by positive torque. We observed previously that R-loop formation times (IN) were dependent on *St3*Cas9 concentration whereas dissociation times (OUT) were not. The same pattern was observed here using the WT and mutant *St3*Cas9 and WT *Sp*Cas9 (Supplementary Figures 2D, 3A and 3B, and Supplementary Table 3). Elevated concentrations of *Sp*Cas9 were required compared to the *St3* enzymes. This difference was variable between *Sp*Cas9 preparations from different sources, most likely due to specific activity variations; subsequent assays (below) used commercial *Sp*Cas9 preparations from New England Biolabs that gave more consistent results.

For all the Cas9s, R-loop IN times were independent of negative torque (Supplementary Figures 2E, 3C and 3D, and Supplementary Table 3); previously observed for WT *St3*Cas9, we suspect this is due to the second order dependence on protein concentration masking the expected torque dependence. The R-loop OUT times showed a dependence on positive torque; dissociation times becomes shorter as positive torque increased (Supplementary Figure 2E). There was little difference between the times observed within experimental error, suggesting that WT *Sp*Cas9 and WT and mutant *St3*Cas9s have similar R-loop stabilities. The RuvC mutant *St3*Cas9(D31A) had a slightly shallower torque dependence, but this is not likely to be significant given experimental uncertainty and DNA-to-DNA variation in the assay.

### 5’ unpaired nucleotides in Streptococcus pyogenes Cas9 gRNA do not have any effects on ribonucleoprotein complex formation

To study the effects of RNA modifications, we first used IVT gRNA with *Sp*Cas9 (Figure 2A, Supplementary Figure 1, Supplementary Tables 1 and 2). Since the number of 5’ guanines can affect transcription efficiency, we produced gRNAs with 1, 2 or 3 unpaired 5’ guanines (1025-G, 1025-GG and 1025-GGG, respectively). The gRNAs target the same target protospacer on pSP1 as in Supplementary Figure 2. In general, sufficient gRNA could be generated by *in vitro* transcription even with a single guanine at the +1 position.

**Figure 2.**
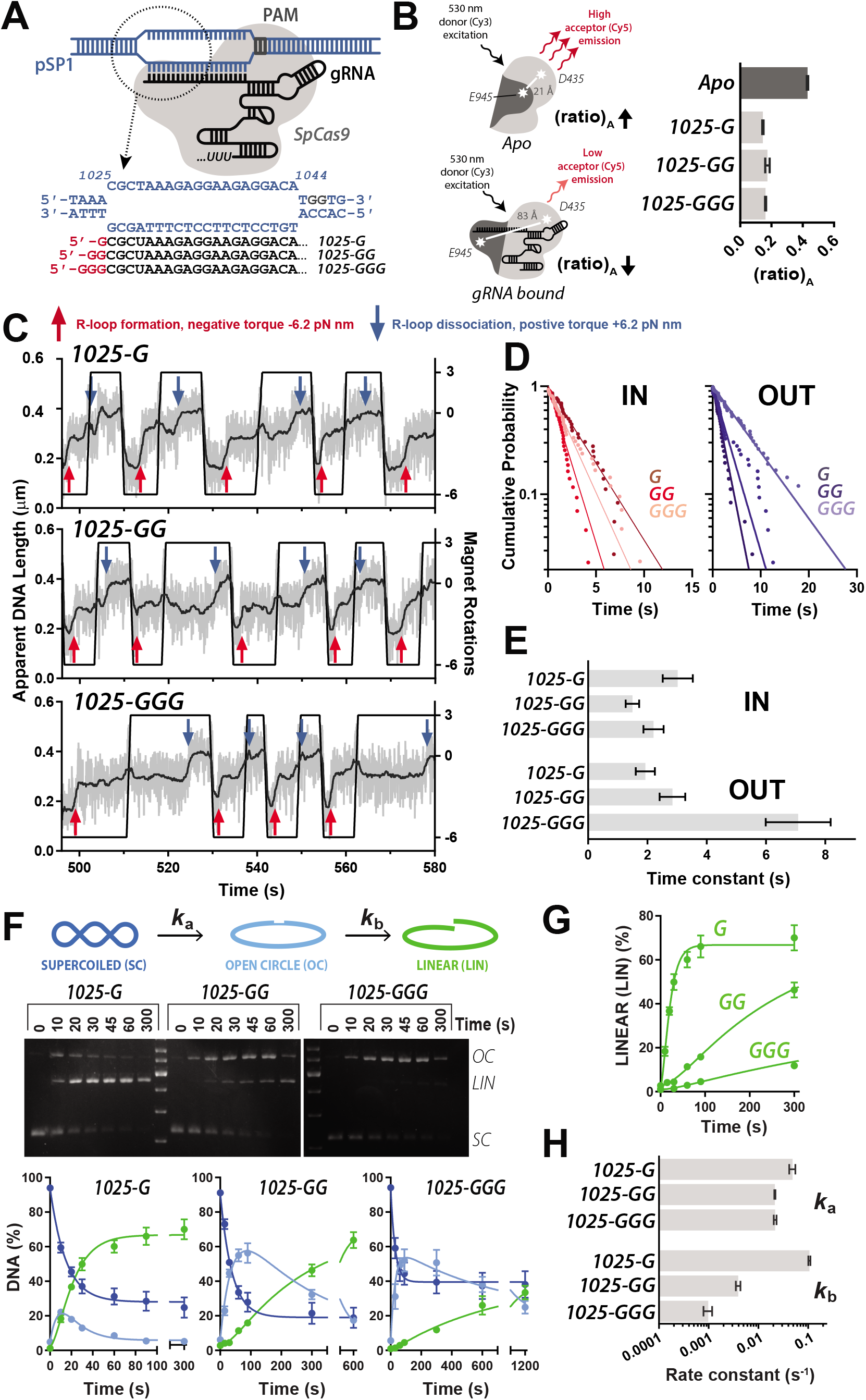
The effects of unpaired guanines at the 5’ end of gRNA on *Streptococcus pyogenes* Cas9. (**A**) Schematic of R-loop formation. *Sp*Cas9 (grey) and the *in vitro* transcribed gRNA (black) binds the PAM (grey) and forms a 20 bp R-loop at the protospacer sequence at 1025-1044 bp of pSP1 (blue). The sequences of the DNA and gRNA spacer sequence are shown. Unpaired 5’ guanines arising from IVT are shown in red. (**B**) FRET-based gRNA loading assay. (*left*) Assay schematic. *Sp*Cas9_hinge_ was randomly labelled at E945C and D435C with Cy3 and Cy5. In the Apo state (*top*), there is high acceptor fluorescence which decreases upon gRNA binding *(bottom)* as fluorophores are moved apart. *(right)* Comparative (ratio)_A_ (*N* = 3 ± SD) for loading gRNAs with 1, 2 or 3 unpaired 5’ guanines (light grey) relative to Apo Cas9 (dark grey). (**C**) Example MT traces of R-loop cycling (at 10 turns s^-1^) to measure R-loop formation (IN, red arrows) and dissociation (OUT, blue arrows). Raw (grey) and 2 Hz smoothed data (black) are shown. Each trace represents measurements on the same single DNA. (**D**) Inverted cumulative probability distributions for the IN and OUT times for each gRNA. Solid lines are single exponential fits. (**E**) Mean IN and OUT times and standard error (*N* = 36 to 43, Supplementary Table 3) from the fits in panel D. (**F**) Supercoiled plasmid cleavage assay. Assay schematic (*top*). The first strand of supercoiled DNA (SC, dark blue) is cleaved to give an open circle (OC, light blue) intermediate, followed by second strand cleavage resulting in linear DNA (LIN, green). These three forms of DNA can be separated by agarose gel electrophoresis *(middle).* Tritiated-DNA bands are cut out from agarose gel and quantified by scintillation counting (*bottom*, *N* = 3 ± SD). The kinetic model (Materials and Methods, Supplementary Figure 6) was simultaneously fitted by numerical integration to each repeat separately, and the kinetic constants averaged (Supplementary Table 4). Solid lines are simulations using the average values. (**G**) Comparison of the appearance of LIN product and fitted profiles for each gRNA taken from panel F. (**H**) Average apparent rate constants for the first and second strand cleavage calculated from the fits in panel F (*N* = 3-4 ± SD) (Supplementary Table 4).

Cas9 loading of gRNAs was followed using a Förster resonance energy transfer (FRET)-based assay developed by Sternberg et al [Materials and Methods, (19,25)]. In the *Sp*Cas9_hinge_ mutant, the natural cysteines are mutated to serine, and positions E945 and D435 are mutated to cysteine for fluorophore labelling (Materials and Methods). In the Apo state, the labelled residues are close (21 Å), resulting in high Cy5 acceptor fluorescence (Figure 2B). As a gRNA is loaded, the labelled residues move ~60 Å, resulting in a corresponding decrease in acceptor fluorescence relative to donor fluorescence. This change scales with increasing molar ratio of gRNA to Cas9 (Supplementary Figure 4A). The ratio between the Cy5 peak as excited by Cy3 fluorescence relative to direct excitation with 630 nm light, “(ratio)_A_” (Supplementary Figures 4B and 4C), was used as a proxy readout for energy transfer (19). In the Apo state, where there is high donor-acceptor energy transfer, (ratio)_A_ was 0.430. Conversely, (ratio)_A_ was lower (0.145-0.173) for each of the three IVT gRNA (Figure 2B). The similar range suggests that there is little or no difference in RNP loading with 1, 2 or 3 additional guanines at the 5’ end of the gRNA.

### 5’ unpaired nucleotides in *Streptococcus pyogenes* Cas9 gRNA have only modest effects on R-loop dissociation kinetics

The three gRNA were then tested in the MT assay using the same DNA molecule. Each RNP tested showed DNA length changes consistent with IN and OUT events as expected for a 20 bp R-loop (Figure 2C). The IN and OUT times and calculated time constants are shown in Figure 2D and 2E, and in Supplementary Table 3. Within error the IN times were similar for the three gRNAs. We cannot directly compare the data to Supplementary Figure 2 as the Cas9 preparations and concentrations were different. The OUT times for 1025-G and 1025-GG were similar to the value observed with *Sp*Cas9 and the crRNA:tracrRNA in Supplementary Figure 2E (the OUT times being independent of concentration). For 1025-GGG, the OUT time was ~two-fold longer. Any rearrangement in RuvC and or in the DNA:RNA hybrid to accommodate 3 unpaired 5’ guanines does not measurably affect the R-loop formation kinetics but does stabilises the structure moderately once formed.

### Unpaired 5’ nucleotides in Streptococcus pyogenes Cas9 gRNA have significant effects on the rates of DNA cleavage

Based on the modest effects on R-loop dynamics observed in Figures 2D and 2E, we expected that nuclease activity would be only moderately affected. DNA cleavage was measured using ^3^H-labelled plasmid DNA separated by agarose gel electrophoresis and the DNA bands quantified by scintillation counting (Materials and Methods) (Figure 2F). Negatively supercoiled DNA has the advantage in nuclease activity measurements that R-loop formation is supported by topology and is unlikely to be rate-limiting as it might be on unconstrained linear DNA (20). Additionally, the nicked intermediate (Open Circle, OC) is readily separated from supercoiled plasmid substrate (SC) and linear product (LIN). The data was fitted to an A-to-B-to-C reaction (Supplementary Figure 6A) to yield apparent rate constants for the first (*k*_a_) and second (*k*_b_) strand cleavages (Materials and Methods, Figure 2H, Supplementary Table 4). The cleavage assays shown in Figure 2F do not go to completion due to low specific activity of the SpCas9 preparation; this does not affect the observed rate constants. More complete DNA cleavage was observed in Figures 3 and 6 with other *Sp*Cas9 preparations. The appearance of the linear product with each of the gRNAs is compared in Figure 2G.

**Figure 3.**
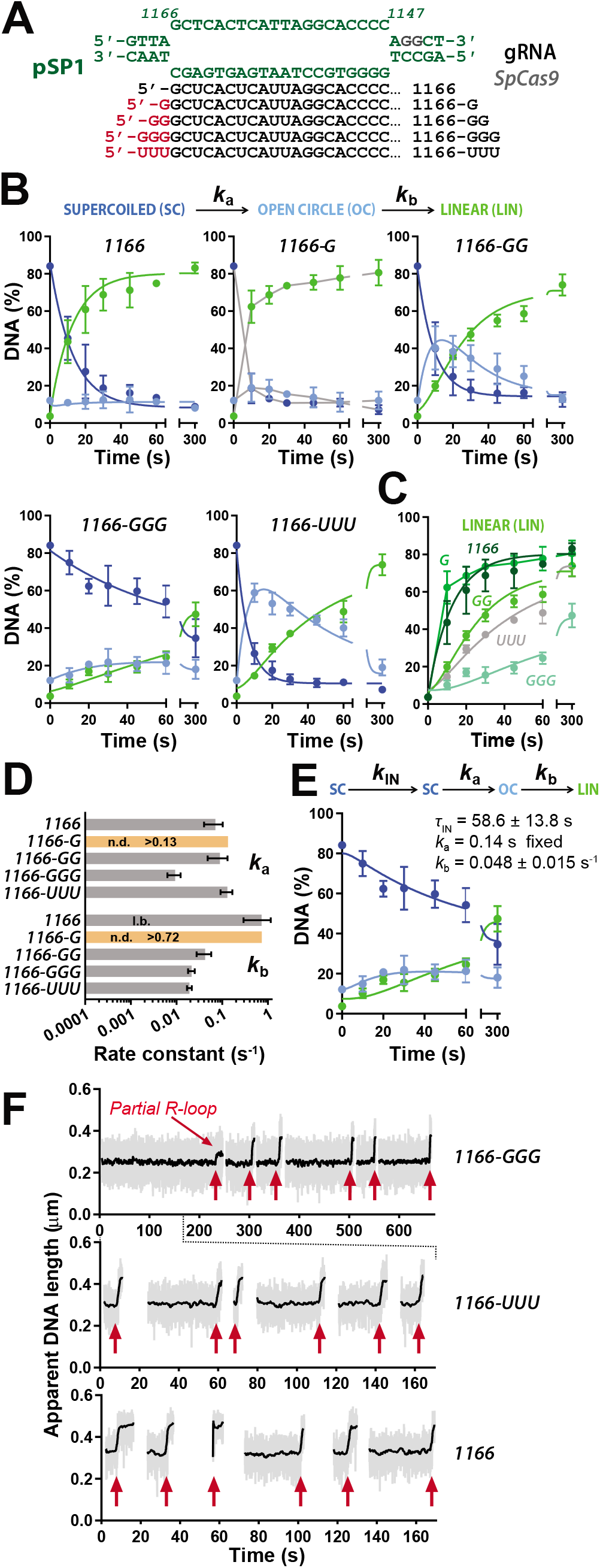
The effects of unpaired bases at the 5’ end of gRNA on *Streptococcus pyogenes* Cas9 at an alternative protospacer sequence. (**A**) Sequence of the 20 bp R-loop at the protospacer sequence at 1166-1147 bp of pSP1 (green). 5’ unpaired bases are shown in red for each of the gRNAs. (**B**) Quantified supercoiled plasmid cleavage assays from gels shown in Supplementary Figure 7A. The kinetic model (Materials and Methods, Supplementary Figure 6) was simultaneously fitted by numerical integration to each repeat separately, and the kinetic constants averaged (Supplementary Table 4). Solid lines are simulations using the average values. (**C**) Comparison of the appearance of LIN product and fitted profiles for each gRNA taken from panel B. (**D**) Average apparent rate constants for the first and second strand cleavage calculated from the fits in panel B (*N* = 3 ± SD) (Supplementary Table 4). (**E**) The alternative kinetic model that incorporates R-loop formation (Supplementary Figure 6) was simultaneously fitted by numerical integration to each repeat separately of the 1166-GGG data. and the kinetic constants averaged (Supplementary Table 4). Solid lines are simulations using the average values. *K*_a_ was fixed at 0.14 s^-1^ based on the average value in panel D. (**F**) Example MT traces showing R-loop formation events (red arrows). Raw (grey) and 2 Hz smoothed data (black) are shown. Each trace represents measurements on the same single DNA.

The rate constants for the first strand cleavage were comparable, with the 1025-GG and 1025-GGG gRNA being only ~2-fold slower than the 1025-G gRNA (Figure 2H, Supplementary Table 4). Surprisingly however, there was a marked difference in the rate of the second strand cleavage (Figure 2G), with the rate constant (*k*_b_) for 1025-GG and 1025-GGG being one or two orders of magnitude slower than 1025-G, respectively (Figure 2H, Supplementary Table 4). The moderate increase in R-loop stability observed in Figure 2E thus corresponds with a slower second strand cleavage. However, the events may not be directly correlated since 1025-GG has largely identical R-loop dynamics to 1025-G yet quite different second strand cleavage kinetics.

In their recent study, Okafor et al suggested that effects of unpaired 5’ nucleotides can be sequence context specific (13). To explore this, we targeted an alternative protospacer sequence on pSP1, where the 20th nucleotide on the protospacer DNA targeted strand is a cytosine that can base pair with a terminal 5’ guanine on the gRNA spacer (Figure 3A). We used synthetic gRNAs rather than IVT gRNAs (Supplementary Table 1), so that we could explore nucleotides other than guanine at the 5’ end. We tested the effect of 0, 1, 2 or 3 unpaired 5’ guanines (1166, 1166-G, 1166-GG and 1166-GGG, respectively), and 3 unpaired 5’ uracils (1166-UUU), to explore whether the inhibitory effect seen in Figures 2F-H was specific to the bulkier purine.

Plasmid cleavage assays using the 1166-family of gRNAs (Supplementary Figure 7A), were quantified and fitted (Figure 3B), with the apparent cleavage rate constants presented in Figure 3D and Supplementary Table 4. The appearance of the linear product with each of the gRNAs is compared in Figure 3C. DNA cleavage with 1166-G was too fast to fit reliably with the model. 1166 (and 1166-G) did not produce any measurable increase in nicked intermediate, due to the second strand cleavage being faster than the first strand cleavage. This difference from the data in Figure 2F highlights that the relative rates of cleavage the first and second strands can vary with protospacer/gRNA sequence.

Apart from 1166-GGG, the apparent rate constants for first strand cleavage were similar (Figure 3D). However, as observed with the 1025 sequence, addition of 2 or more unpaired 5’ nucleotides resulted in >10-fold slower apparent rates constants for second strand cleavage relative to the 1166 gRNA. The addition of 3 unpaired 5’ uracils was worse than adding 2 unpaired 5’ guanines, suggesting that the inhibitory effect on cleavage is not specific to guanines and scales with length. However, the effect of 3 unpaired 5’ guanine residues was markedly worse in terms of the apparent appearance of linear DNA (Figure 3C); the fitted kinetic values suggest this is because the first strand cleavage was also slow (Figure 3D, Supplementary Table 4).

An alternative explanation for the cleavage profile with the 1166-GGG gRNA in Figure 3B is that the apparent cleavage rate constants are being influenced by slower R-loop formation kinetics using this gRNA compared to the others. A rate limiting R-loop formation step can mask subsequent DNA cleavage steps (20). To consider this, the 1166-GGG data was re-fitted with a model that incorporated R-loop formation as a one-step process (*k*_I_N) prior to first strand cleavage (Supplementary Figure 6B). Where *k*_IN_ is rate-limiting, a fitted value for *k*_a_ cannot be returned based on the cleavage data alone; *k*_a_ was therefore fixed as the average value for the other gRNA (0.14 s^-1^), and *k*_IN_ and *k*b were allowed to float. This model could also describe the data (Figure 3E), with an apparent *k*_b_ similar to that for 1166-GG but with a slow apparent R-loop formation.

To test whether the 3 unpaired 5’ guanines on 1166-GGG gRNA were changing the R-loop formation kinetics, we repeated the MT assay using 1166-GGG, 1166-UUU and 1166 gRNAs (Figure 3F, only ON events shown). R-loop formation events were markedly slower using 1166-GGG. Note that we cannot directly compare R-loop formation times in the tweezers to the apparent *k*_IN_ value from the DNA cleavage assays as the *Sp*Cas9 RNP concentrations are different and R-loop formation has a second-order component (Supplementary Figure 2).

### Slow second strand cleavage is due to the influence of the unpaired 5’ nucleotides on the activity of the RuvC nuclease domain

Okafor et al. (13) proposed that due to close interaction between the RuvC nuclease domain and the 5’ end of the spacer RNA, unpaired residues may alter Cas9 activity. The slower rate of second strand cleavage observed here with both the 1025 and 1166 gRNAs (Figures 2 and 3) could therefore be due to misalignment of the RuvC nuclease that slows cleavage of the non-targeted strand. To explore this, we compared the DNA cleavage profiles using *st3*Cas9, *st3*Cas9(D31A) and *st3*Cas9(N891A) with the *St3*-specific 1025-G and 1025-GGG gRNAs (Figure 4, Supplementary Figures 1 and 7B, Supplementary Table 1).

**Figure 4.**
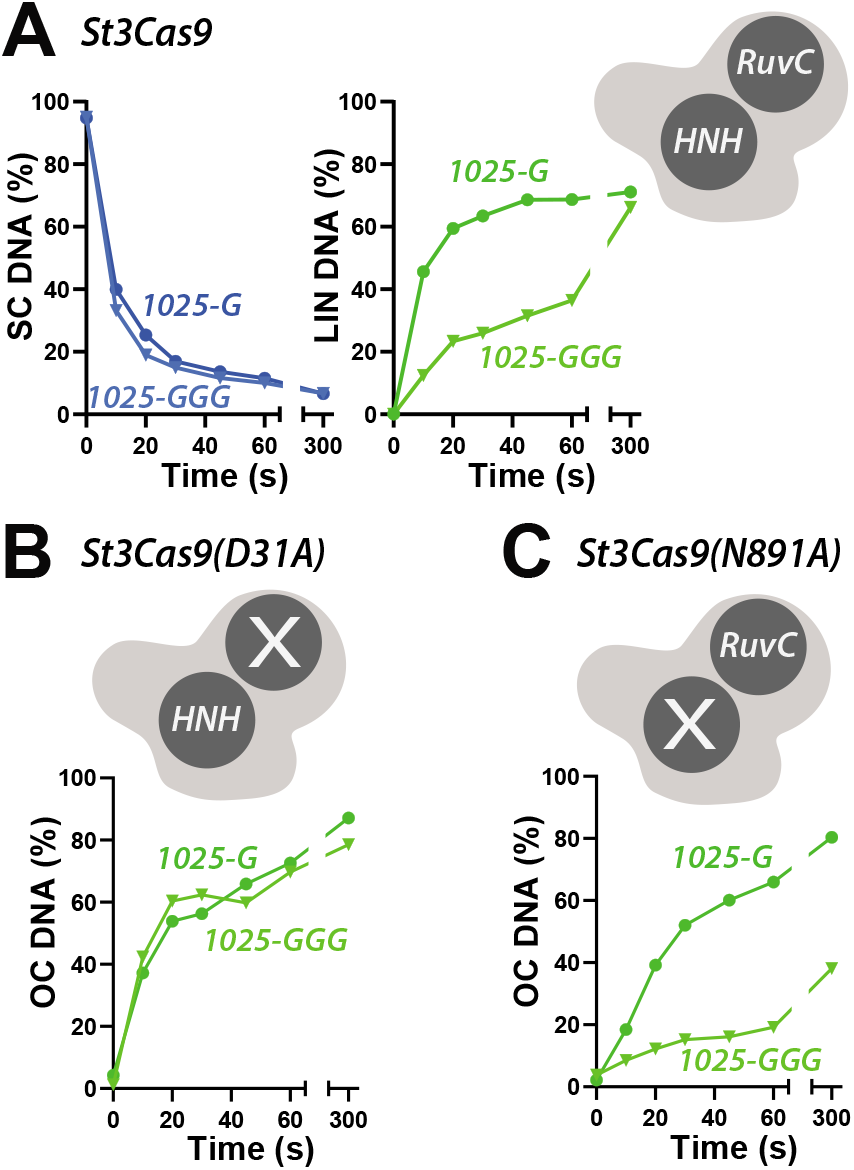
Unpaired bases at the 5’ end of gRNA slow the rate of DNA cleavage by the RuvC nuclease domain but not by the HNH nuclease domain. Quantified supercoiled plasmid cleavage assays from gels shown in Supplementary Figure 7B using *Streptococcus thermophilus* DGCC7710 CRISPR3 Cas9 WT (**A**), RuvC mutant D31A (**B**), or HNH mutant N891A (**C**). For clarity, only the disappearance of supercoiled (SC) DNA and appearance of linear (LIN) product for WT are compared to the appearance of nicked (OC) product for the nicking mutants.

As observed with *Sp*Cas9, the first strand cleavage by *St*Cas9 was unaffected by the number of unpaired 5’ guanines but there was an inhibition of second strand cleavage with 1025-GGG relative to 1025-G (Figure 4A). For *St3*Cas9(D31A), where only the HNH nuclease is active, the rate of appearance of nicked product was similar with either gRNA (Figure 4B). In contrast, for *St3*Cas9(N891A), where only the RuvC nuclease is active, there was a marked inhibition of the rate of appearance of nicked product using 1025-GGG. This result is consistent with the slower second strand cleavage being due to disruption of docking of the RuvC nuclease due to the 3 unpaired guanine residues. We note that we cannot distinguish based on our data between an ordered strand cleavage model (HNH cutting the targeted strand first, followed by RuvC cutting the non-targeted strand second) and a random cleavage model (20).

### A structured RNA hairpin a the 5’ end of a gRNA can produce shorter hyperstable R-loops that do not support DNA cleavage

To explore the effect of adding a structured RNA aptamer to *Sp*Cas9 *g*RNA, we utilised a 20 bp RP hairpin (Figure 5A) derived from a stem loop structure in the nuclear RNaseP RNP, that can direct import of RNA into mitochondria (21,26). A library of gRNAs was designed in which the RP hairpin was added at one of four gRNA locations (Figure 5A, Supplementary Figure 1): the 5’ end (5’ RP); the Upper Stem (RP US); two alternative positions in the first Hairpin (RP H1 and RP H2); and the 3’ end (3’ RP). gRNAs were synthesised by IVT (Supplementary Table 2); this incorporated a single unpaired 5’ guanine in all gRNAs and a variable run of uracils at the 3’ end. We chose to target the 1025 protospacer sequence in pSP1 as in Figures 2 and 4.

**Figure 5.**
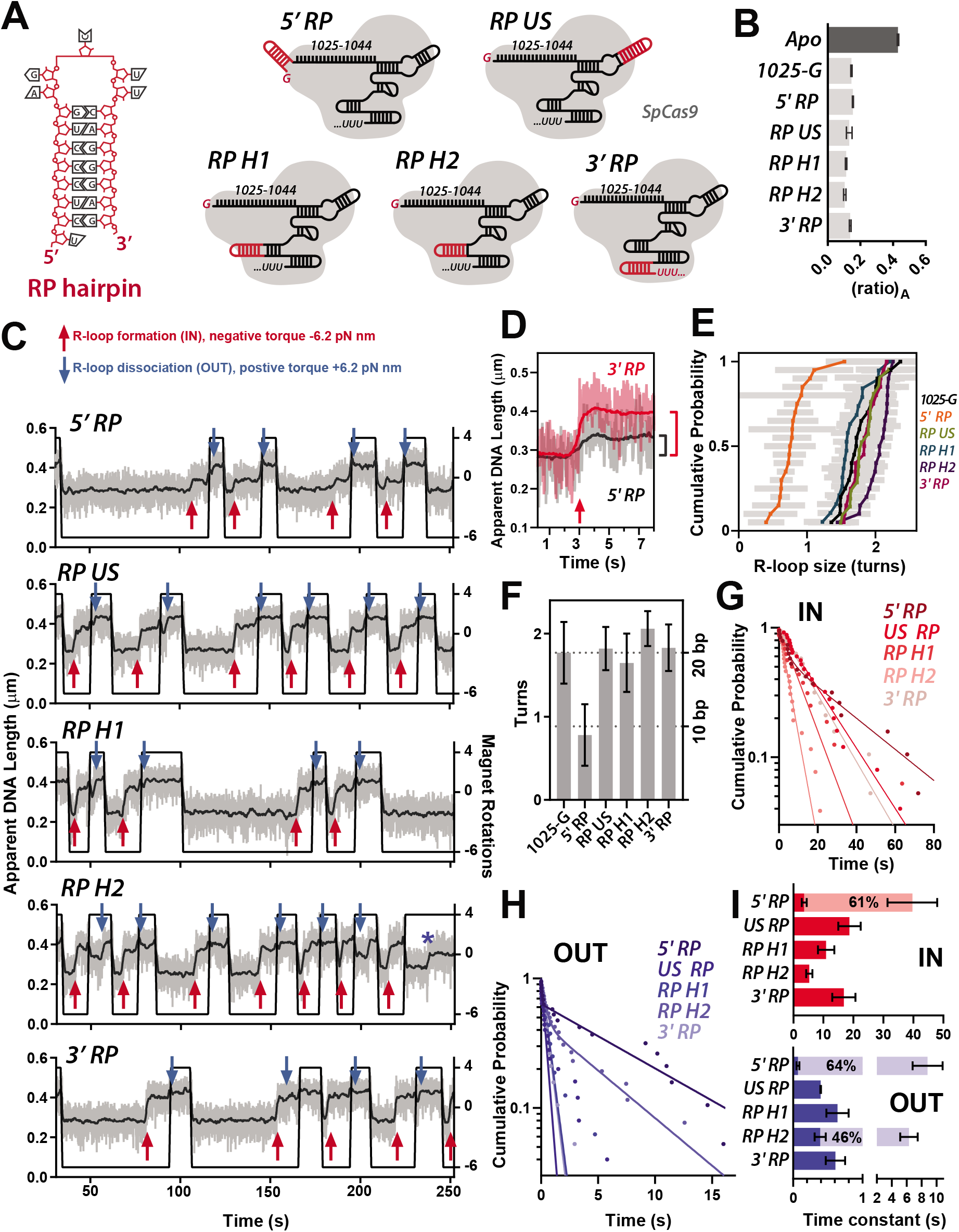
The effect of the RP RNA hairpin on R-loop dynamics. (**A**) Sequence of the RP hairpin and schematics of *Streptococcus pyogenes* Cas9 (grey) and the *in vitro* transcribed gRNAs (black) (sequences in Supplementary Figure 1). (**B**) FRET-based gRNA loading assay. Comparative (ratio)_A_ for loading gRNAs as shown (light grey) relative to Apo Cas9 (dark grey) *(N* = 3 ± SD). (**C**) Example MT traces of R-loop cycling (at 10 turns s^-1^) to measure R-loop formation (IN, red arrows) and dissociation (OUT, blue arrows). Raw (grey) and 2 Hz smoothed data (black) are shown. (**D**) Comparison of an R-loop formation event with 3’ RP (red) and 3’ RP (black) gRNA. (**E**) cumulative probability of R-loop size in turns estimated from R-loop cycling experiments in Supplementary Figure 8 (Materials and Methods). Grey bars are the standard error for each estimation. (**F**) Average R-loop size in turns (± SD) for each of the gRNAs. The average turns for 1025-G, RP US, RP H1, RP H2 and 3’ RP was assumed to estimate a full length R-loop and used to set R-loop size in bp. (**G, H**) Inverted cumulative probability distributions for the IN and OUT times for each gRNA. Solid lines are single exponential fits or double exponential fits (5’ RP IN; 5’ RP and RP H2 for OUT). (**I**) Mean IN and OUT times and standard error (*N* = 19 to 43, Supplementary Table 3) from the fits in panel D. For 5’ RP IN events, and 5’ RP and RP H2 OUT events, the light and dark bars are the two constants from a double exponential fit with the percentage shown for the amplitude of the slower events.

The loading of the modified gRNA library and *Sp*Cas9 into RNPs was measured using the FRET-based *Sp*Cas9_hinge_ assay described above (19), and compared to 1025-G (Figure 5B, Supplementary Figure 5). The gRNA RP library gave similar relative acceptor fluorescence changes in the (ratio)_A_ range 0.104-0.154, comparable to the value seen with 1025-G (0.145). We conclude that RP modifications do not have a significant impact on the Cas9 structure with respect to REC lobe positioning during gRNA loading.

The MT assay was used to assess the effect of the RP hairpin on R-loop formation and dissociation (Figure 5C). In all cases, stable R-loops were observed that could be reversed by application of positive torque. It was noticeable that the change in bead height produced by the 5’ RP gRNA was smaller than the characteristic 20 bp R-loop change with the other gRNAs; R-loop events from 5’ RP and 3’ RP are compared in Figure 5D. Rotation curves in the absence and presence of *Sp*Cas9 were used to quantify the shift in the number of DNA turns that are captured by R-loop formation (Supplementary Figure 8, Figure 5E, F) (20). The 5’ RP gRNA produced an R-loop that captured only ~0.78 DNA turns compared to the average values for the other gRNAs of 1.65 – 2.06 turns. This suggests the 5’ RP gRNA only supports formation of a stable ~9 bp R-loop, a size which corresponds to that expected for the seed region. It is also possible that RP H2 supports a slightly larger R-loop (Figure 5E,F), but the data is close to the error limits of R-loop size estimation and was collected on a different DNA molecule to the other gRNAs.

The R-loop IN and OUT times were calculated at single negative and positive torque values (Figure 5 G-I, Supplementary Table 3). The IN times were all relatively slow; for 5’ RP the data had a double exponential distribution, with ~61% of events being very slow (~36 s) and 39% of events being markedly faster (~3.5 s). The OUT times for US RP, H1 RP and 3’ RP were all faster than 1025-G, indicating a moderate loss of R-loop stability. For both 5’ RP and H2 RP, the OUT times were biphasic, with a proportion of events being slower to dissociate. These may indicate abnormal R-loop structures that are hyperstablised.

Given the alterations to the R-loop dynamics and size, we expected that DNA cleavage kinetics would also be alerted for at least some of these modified gRNAs. However, for RP US, RP H1, RP H2 and 3’ RP, the DNA cleavage profiles and the apparent rate constants for the first and second strand cleavages were very similar to 1025-G (Figures 6A and 6C, Supplementary Figure 7C, Supplementary Table 4). In comparison, only a low percentage of DNA nicking was observed over 16 hours using 5’ RP gRNA (Figures 6B and 6C). We reasoned that the inhibition by RP may be because the structured hairpin is too close to the DNA:RNA hybrid. However, adding uracil residues between the RP hairpin and spacer sequence to give more flexibility (Supplementary Table 2), did not activate DNA cleavage to measurable levels (Figure 6B), although the 5’ RP U4 gRNA did support formation of a small amount of linear DNA after 16 hours incubation. Despite being relatively stable (Figure 5I), the shorter R-loop produced by the 5’ RP gRNA cannot activate DNA cleavage over a reasonable timescale.

**Figure 6.**
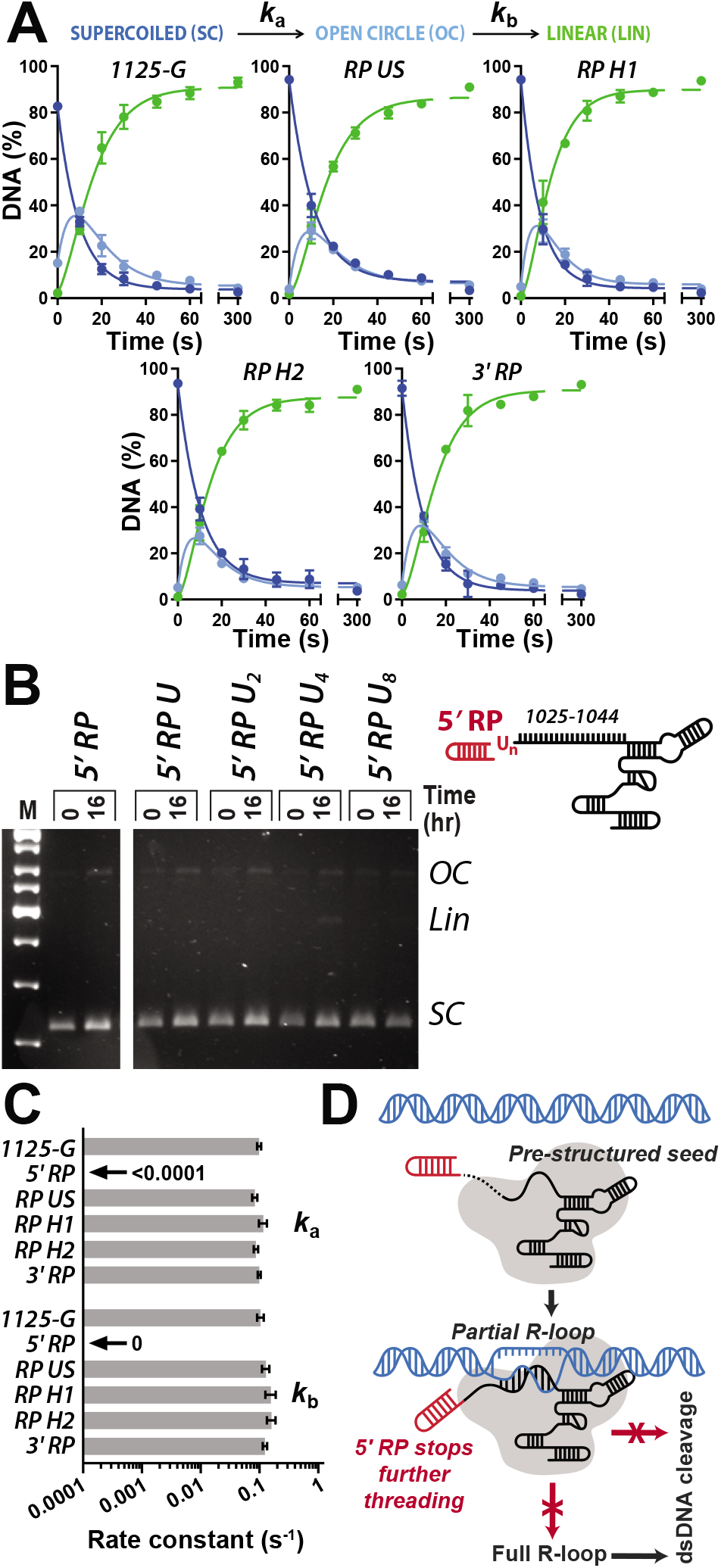
5’ RP modification of gRNA blocks DNA cleavage by *Streptococcus pyogenes* Cas9. (**A**) Quantified supercoiled plasmid cleavage assays from gels shown in Supplementary Figure 6C. The kinetic model (Materials and Methods, Supplementary Figure 6) was simultaneously fitted by numerical integration to each repeat separately, and the kinetic constants averaged (Supplementary Table 4). Solid lines are simulations using the average values. (**B**) Cartoon of the 5’ RP gRNAs modified to include 1, 2, 4 or 8 uracils. Agarose gel of pSP1 cleavage reactions using *Sp*Cas9 and the gRNAs, as indicated. (**C**) Average apparent rate constants for the first and second strand cleavage calculated from the fits in panel A (*N* = 3 ± SD) (Supplementary Table 4) or estimated from panel B. (**D**) Model for partial R-loop formation by 5’ RP gRNA. See text for further details.

## DISCUSSION

gRNA engineering allows a wide range of enhancements to CRISPR-Cas activity (7). Chemical synthesis of gRNAs has become routine, and modification of nucleotides and addition of functional groups have increased nuclease-resistance and/or enhanced nuclear localization. Additionally, fusion of additional RNA sequences/structures to gRNA has been exploited to add new functionality to CRISPR-Cas. However, our single-molecule and ensemble measurements of gRNA loading, R-loop formation and DNA cleavage indicate that modified gRNAs can have significant inhibitory effects on DNA cleavage in a target sequence-dependent manner, and can influence the size and stability of the R-loop. These possible changes in CRISPR-Cas activity need to be considered when testing any new designs.

The 5’ modification of Cas9 gRNA with a 20 nt RP hairpin had the striking effect of preventing DNA cleavage due, most likely, to the formation of an incomplete R-loop corresponding to the seed sequence. Previous single molecule data suggested a critical cut off in R-loop length at around 10-11 bp below which the R-loop could not form and above which R-loop stability was similar (18). Consistent with the prevailing model for activation of the Cas9 nuclease activity (19,27), this size of RNA:DNA hybrid will not engage the HNH domain. Very low levels of DNA cleavage were only observed here over very long timescales; this may reflect non-specific cleavage activity or a very slow formation of a full-length R-loop that was not observed in the MT experiments.

In contrast, fusion of the RP hairpin at the Upper Stem, Hairpin and 3’ end had much less drastic effects on R-loop formation and no measurable effect on DNA cleavage. These observations highlight that modifications to other parts of the gRNA will be more likely to be successful. Consistent with this, these gRNA features have been shown to accommodate RNA aptamer scaffolds with a range of sequences and sizes, e.g. (15,28–31). Often gRNA fusions are used to provide a binding sequence to colocalise a protein or fluorescent label and are hence used with dCas9 rather than nuclease active WT Cas9. Thus, DNA cleavage activity is not always assessed, and one cannot rule out that atypical R-loop structures may be produced.

A possible explanation for the abridged R-loop zipping using 5’ RP is suggested in Figure 6D. Following the formation of the DNA:RNA hybrid in the seed region, the 5’ end of the gRNA may need to thread around the targeted DNA strand. This threading may be inhibited by the addition of a bulky structured RNA. Other studies have successfully modified the 5’ end of gRNA and retained cleavage activity (a proxy for complete R-loop formation), e.g. (16,17,32). However, in many cases the fusions were not highly structured. The type I Cascade complex can thread a 3’ crRNA hairpin when forming its 33 bp R-loop (18,33), but this is a quite different structural complex to Cas9. It may be that the effect seen here is unique to the RP hairpin and that empirical design can be used to select for RNA structures that have the desired effect and to avoid unwanted ones. There may also be protospacer specific effects, as observed for the short 5’ modifications (13). The unusual activity of 5’ RP gRNA could be exploited to form stable R-loops without activating DNA cleavage. Multiplex expression of unmodified and modified gRNAs could allow for simultaneous site-specific editing and site-specific binding at different locations by WT Cas9.

The more surprising observation from our study was that two or more unpaired nucleotides at the 5’ end of Cas9 gRNAs can have significant effects on the activity of the RuvC nuclease domain while, in some cases, having only relatively modest effects on R-loop stability. As noted by Okafor et al. (13), because there are close interactions of the docked RuvC domain with the 5’ end of the RNA observed in crystal structures (9), 5’ modifications may result in rearrangements in the RuvC domain that slows, but does not completely abrogate, the cleavage of the second strand. Consistent with this, we observed that DNA nicking by the RuvC nuclease was affected by 3 unpaired 5’ guanines whereas DNA nicking by the HNH nuclease was not.

Our observations highlight a potential inefficiency of using gRNA with more than one unpaired nucleotide at the 5’ end if the DNA cleavage activity of Cas9 is critical to the application. However, the extent of the effect varied with DNA/RNA sequence, as was also noted by Okafor et al (13). For example, the addition of 3 unpaired 5’ guanines to the 1025 gRNA slowed the second strand cleavage rate, while the same addition to the 1166 gRNA slowed both R-loop formation and second strand cleavage. Additionally, in the single molecule traces with 1166-GGG we also observed occasional events that were smaller than expected for a 20 bp R-loop (partial R-loop marked in Figure 3F). This suggest that even small 5’ modifications can, in some sequence contexts, inhibit the completion of R-loop zipping possibly for the same reason as shown in Figure 6D.

The undesirable effects of unpaired 5’ nucleotides can be avoided. IVT is still relatively efficient with a single guanine which had minimal effect on cleavage efficiency. Similarly, the eukaryotic U6 promoters typically exploited when expressing gRNAs from a gene in cell culture only add a single unpaired nucleotide. By careful selection of protospacer sequences and selection of sites using online tools (34), the base can be incorporated as part of the spacer RNA:DNA hybrid. Alternatively, a hammerhead ribozyme domain can be added to the 5’ end which can remove the mismatched guanine prior to RNP formation and CRISPR activity (12). For RNP transfection-based methods, the price of commercial RNA synthesis using phosphoramidites has decreased significantly in recent years and unmatched 5’ guanines can be excluded from the design.

## Supporting information

Supplementary results and tables

## ACKNOWLDEGEMENTS

We thank Ciaran Guy, Carlos Martinez Santiago and Jia Qi Cheng Zhang for help with experiments and data analysis.

