## Supplementary results and tables for "5’ modifications to CRISPR Cas9 gRNA can change the dynamics and size of R-loops and inhibit DNA cleavage"

[4]CasZyme, Saulėtekio al. 7A, Vilnius, Lithuania

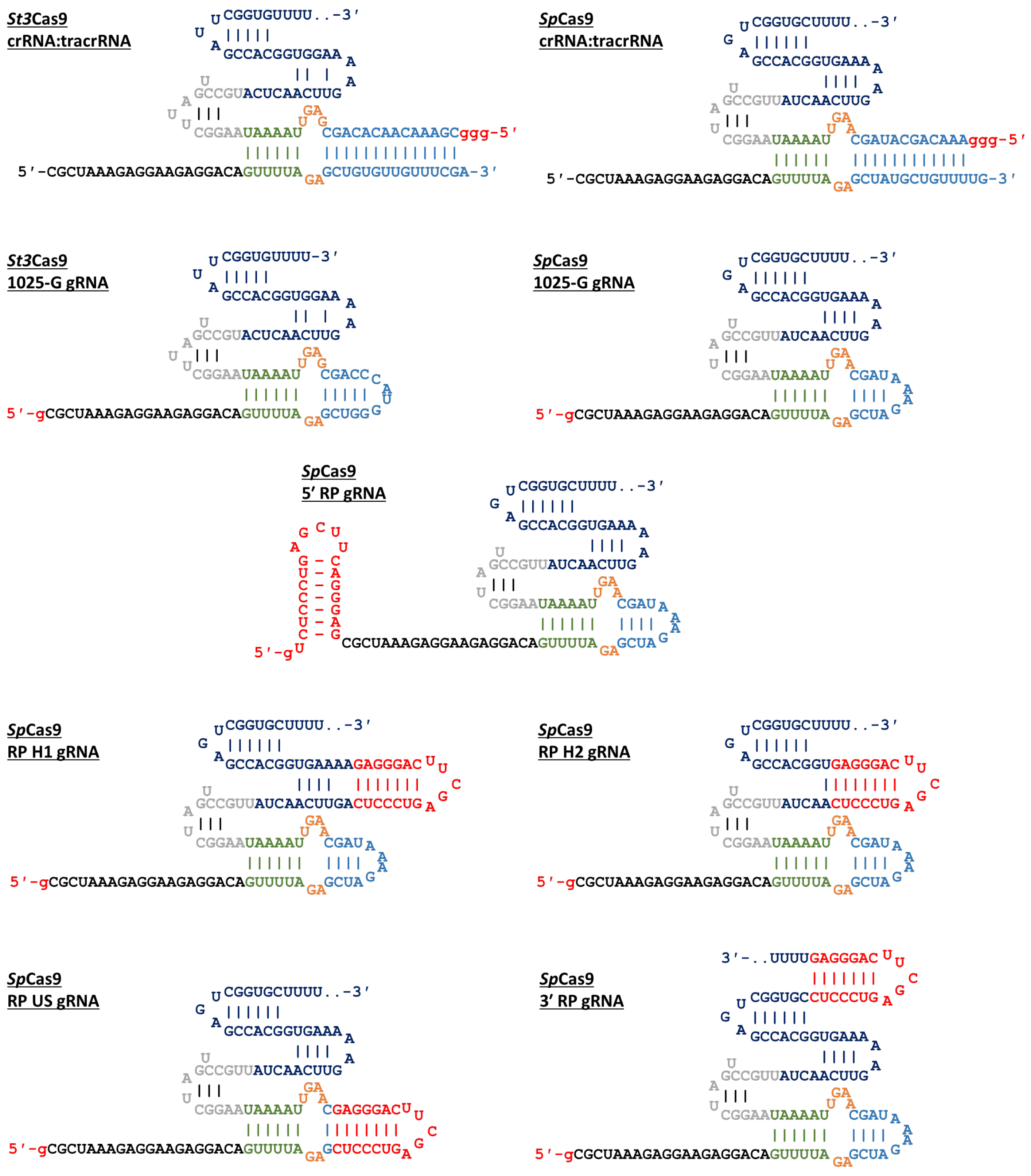


**Supplementary Figure 1.** Sequences and putative folded structure of crRNA:tracrRNA and gRNA used in this study. Sequences are coloured: red, structural additions; black, Spacer; green, Lower Stem; orange, Bulge; light blue, Upper Stem; Grey, Nexus; dark blue, Hairpins. Red guanines at the 5' ends are due to *in vitro* transcription. Dotted lines at 3' ends represent potential variability in the number of uracils due to termination of *in vitro* transcription.

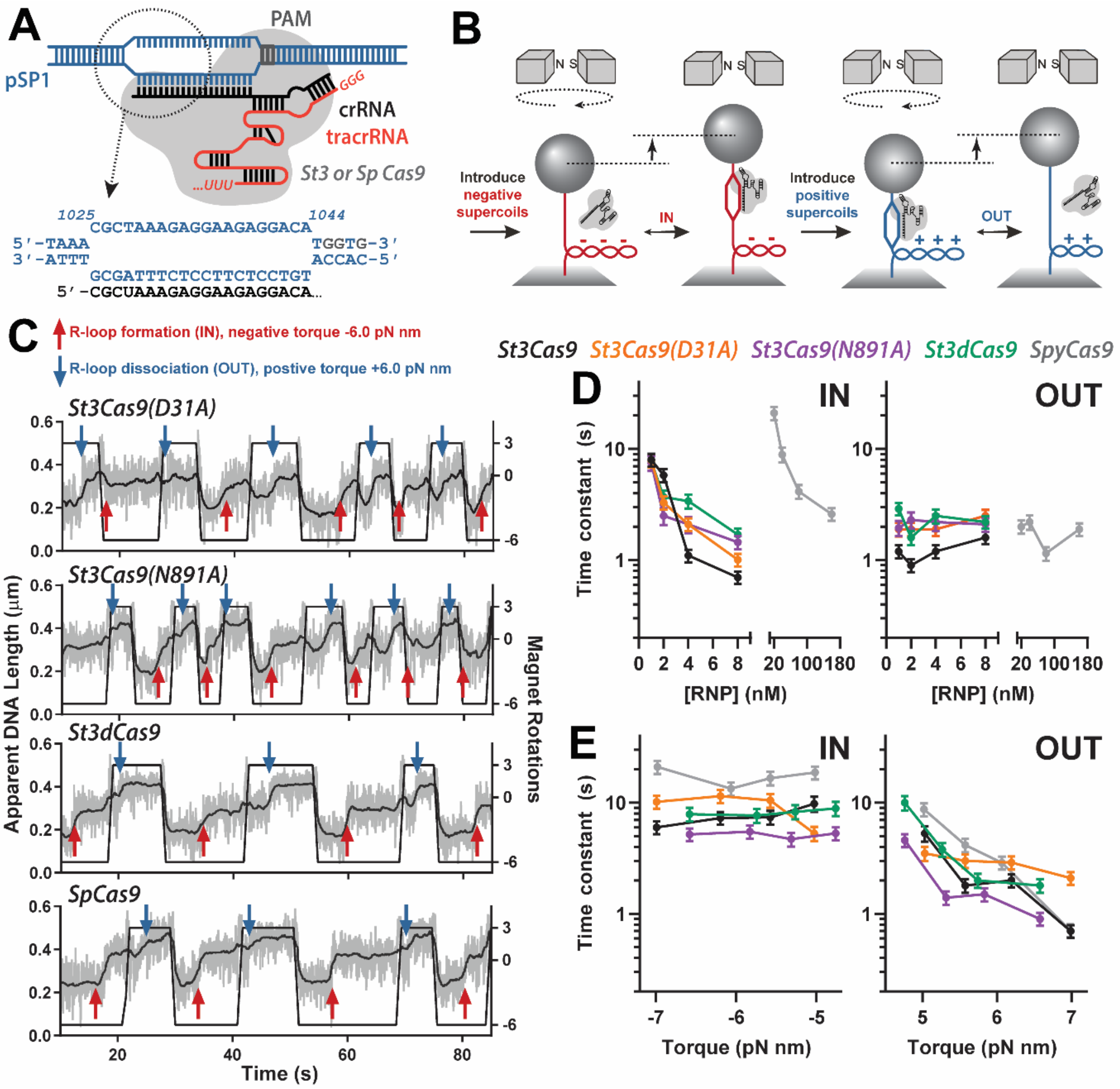


**Supplementary Figure 2.** R-loop dynamics for *Streptococcus pyogenes* and *Streptococcus thermophilus* DGCC7710 CRISPR3 Cas9 with their respective crRNA:tracrRNA. (B) Schematic of R-loop formation. Cas9 (grey) and the cRNA: tracrRNA (black:red) binds the PAM (grey) and forms a 20 bp R-loop at the protospacer sequence at 1025-1044 bp of pSP1 (blue). The sequences of the DNA and crRNA spacer sequence are shown. (**B**) Schematic of the magnetic tweezers (MT) assay. See main text for details. (**C**) Example MT traces of R-loop cycling (at 10 turns $s^{-1}$) to measure R-loop formation (IN, red arrows) and dissociation (OUT, blue arrows). Raw (grey) and 2 Hz smoothed data (black) are shown. Each enzyme trace represents a different single DNA. (**D**) Mean IN and OUT times and standard error (*N* = 31 to 64, Supplementary Table 3) as a function of RNP concentration for WT *St3*Cas9 (black), *St3*Cas9(D31A) (orange), *St3*Cas9(N891A) (purple), *St3*dCas9 (green) and *Sp*Cas9 (grey). (**E**) Mean IN and OUT times and standard error (*N* = 40 to 60, Supplementary Table 3) as a function of torque (enzyme colours as in panel D) using 1 nM RNP for all *Streptococcus thermophilus* RNPs and 10 nM for *Sp*Cas9 RNP. Cumulative probability distributions used to calculate the times in panels D and E are shown in Supplementary Figure 3.

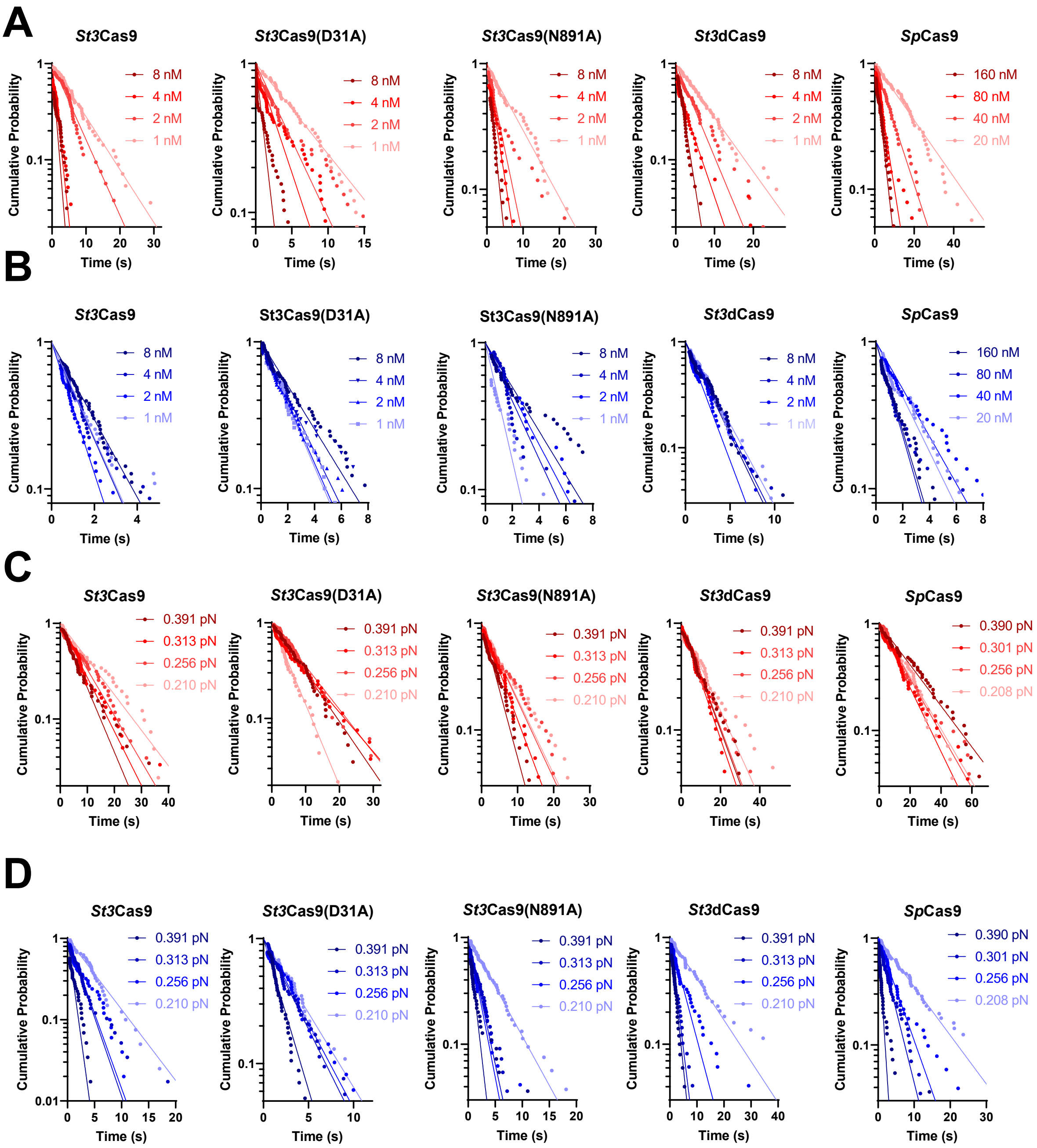


**Supplementary Figure 3.** Cumulative probability distributions for IN and Out events with Cas9 and crRNA:tracrRNA. R-loop cycling in a magnetic tweezers assay (Figure 2B) was used to analyse the R-loop dynamics for WT *St3*Cas9, the nicking mutants *St3*Cas9(D31A) and *St3*Cas9(N891A), the nuclease dead mutant *St3*dCas9, and WT *Sp*Cas9 (Figure 2C). Data were fitted (solid lines) to a single exponential decay apart from those stated in Supplementary Table 3. (**A, B**) R-loop formation (IN, panel A) and dissociation (OUT, panel B) times as a function RNP concentration (Cas9:crRNA:tracrRNA = 1:1:1) at 0.313 pN. (**C, D**) R-loop formation (IN, panel C) and dissociation (OUT, panel D) times as a function of applied DNA stretching force with an RNP concentration of 1 nM for *St3*Cas9 variants and 10 nM for *Sp*Cas9 (with Cas9:crRNA:tracrRNA = 1:1:1).

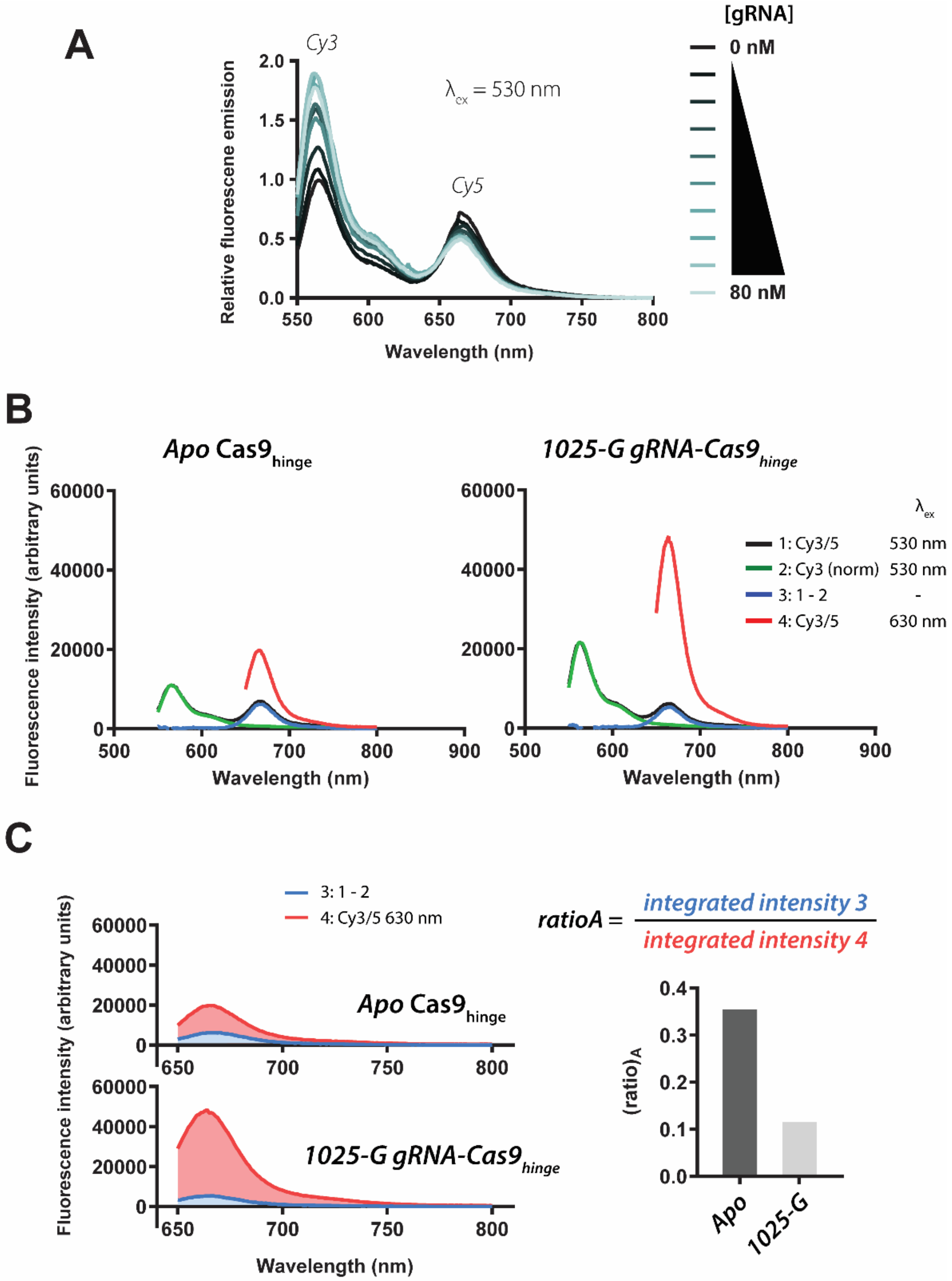


**Supplementary Figure 4.** FRET measurements of gRNA loading with *Sp*Cas9$_{hinge}$. (**A**) *Sp*Cas9$_{hinge}$ was labelled with Cy3, or with Cy3 and Cy5, at positions 435 and 945 (Materials and Methods). In the Apo state of *Sp*Cas9$_{hinge}$-Cy3/5, there is high acceptor fluorescence. Donor (Cy3) fluorescence rises and acceptor (Cy5) fluorescence falls with increasing [gRNA], when the donor fluorophore is excited at 530 nm. [*Sp*Cas9$_{hinge}$] = 50 nM. (**B, C**) (Ratio)$_A$ can be calculated by dividing the integrated intensity between 650 and 800 nm (scan 3) [calculated as the difference between *Sp*Cas9$_{hinge}$-Cy3/5 (scan 1) and *Sp*Cas9$_{hinge}$-Cy3 (scan 2), excited at 530 nm], by the integrated intensity 4 between 650 and 800 nm (scan 4) [calculated from *Sp*Cas9$_{hinge}$-Cy3/5 directly excited at 630 nm]. Example scans are shown for Apo and 1025-G gRNA bound states.

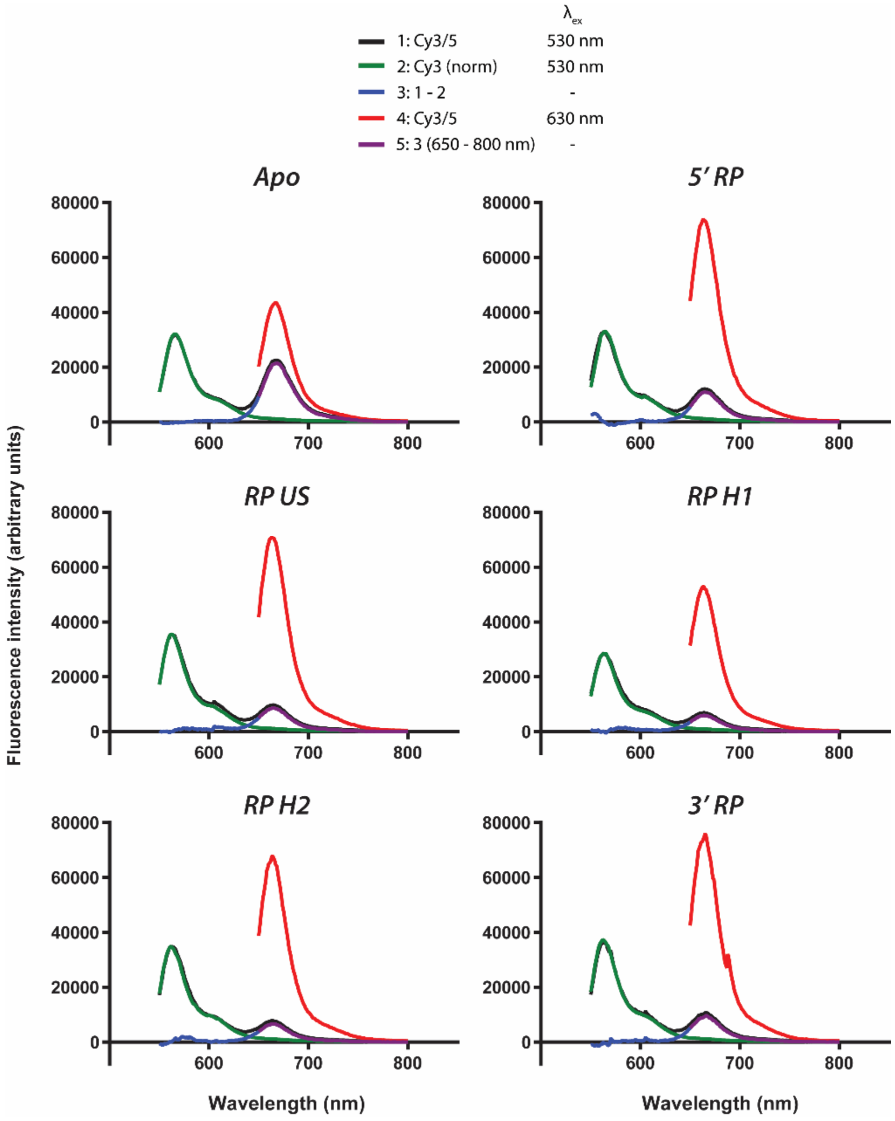


**Supplementary Figure 5.** Example FRET measurements of loading of *Sp*Cas9$_{hinge}$ with RP-modified gRNAs as indicated.

**A**

```
Method RK4
STARTTIME = 0
STOPTIME = 300
DT = 0.02
d/dt (CCC) = -ka*CCC - kini*CCC
d/dt (ucCCC) = kini*CCC
d/dt (OC) = ka*CCC - kb*OC - kini2*OC -ka*OC1
d/dt (OC1) = -ka*OC1 - kini*OC1
d/dt (ucOC) = kini2*OC + kini*OC1
d/dt (LIN) = kbOC
Tot CCC = CCC + ucCCC
Tot OC = OC + OC1 + ucOC
Init CCC = 95
init LIN = 1
init OC1 = 4
init OC = 0
init ucCCC = 0
init ucOC = 0
ka = 0.10
kb = 0.10
kini = 0.02
kini2 = 0.02
```

these values changed to actual values for each run

uncut OC/CCC allows for reaction not going to completion.

these values allowed to be fit by software

**B**

```
METHOD RK4
STARTTIME = 0
STOPTIME = 300
DT = 0.02
d/dt (CCCC)= -kIN*CCCC
d/dt (CCC) = kIN*CCCC - ka*CCC - kini*CCC
d/dt (ucCCC) = kini*CCC
d/dt (OCC1) = -kIN*OCC1
d/dt (OC1) = kIN*OCC1 - ka*OC1 - kini*OC1
d/dt (OC) = ka*CCC + ka*OC1 - kb*OC - kini2*OC
d/dt (ucOC) = kini2*OC + kini*OC1
d/dt (LIN) = kb*OC
TotCCC = CCCC + CCC + ucCCC
TotOC = OCC1 + OC1 + OC + ucOC
init CCCC = 95
init CCC = 0
init LIN = 1
init OCC1 = 4
init OC1 = 0
init OC = 0
init ucCCC = 0
init ucOC = 0
ka = 0.14
kb = 0.10
kini = 0.02
kini2 = 0.02
kIN = 0.01
```

these values changed to actual values for each run

uncut OC/CCC allows for reaction not going to completion.

these values allowed to be fit by the software

**Supplementary Figure 6.** DNA cleavage models used for numerical integration with Berkeley Madonna. (**A**) Standard A-B-C dsDNA cleavage model that allows for incomplete cleavage. (**B**) Modified A’-A-B-C dsDNA cleavage model that allows for an R-loop formation step prior to cleavage, and for incomplete cleavage.

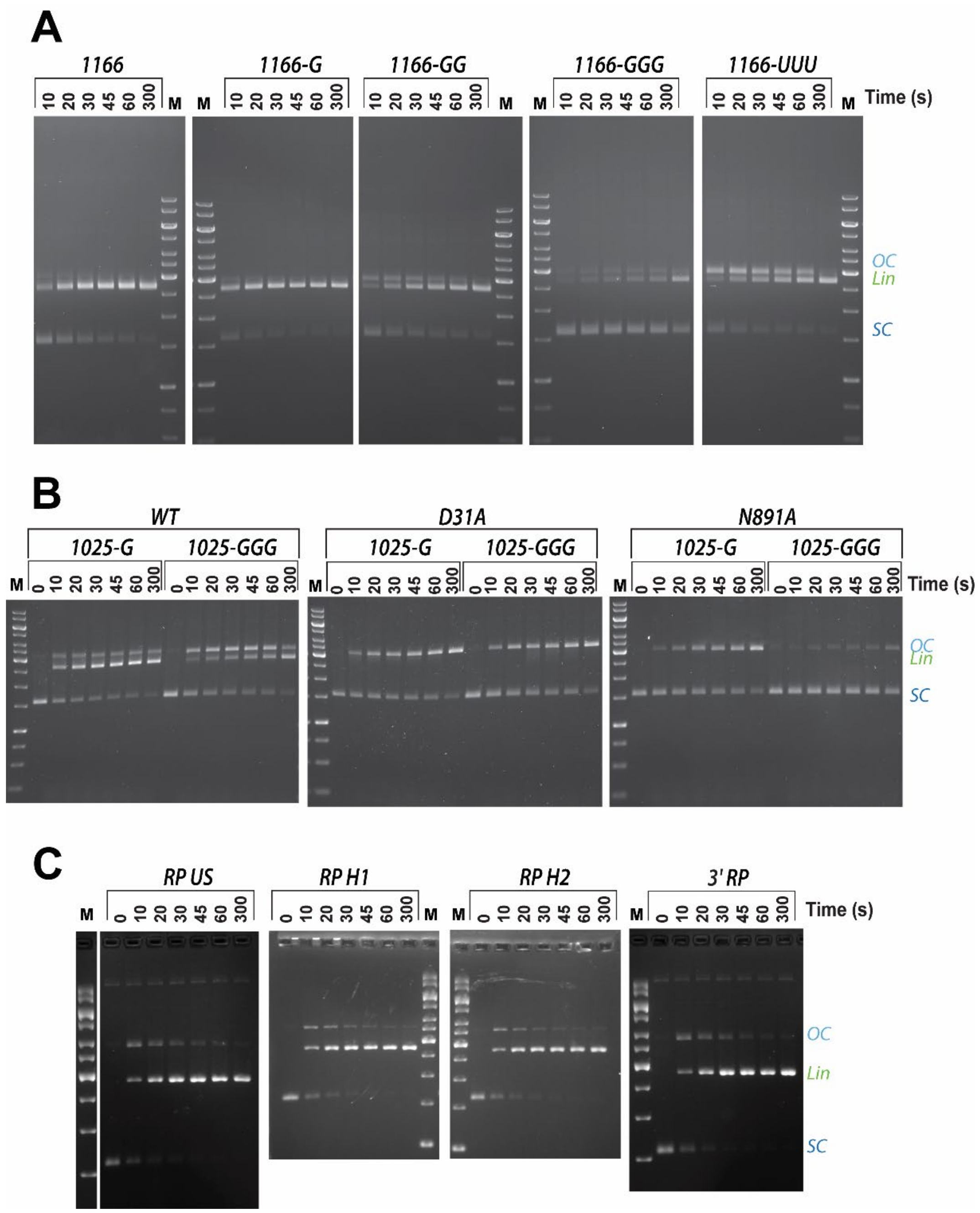


**Supplementary Figure 7.** Example agarose gels of DNA cleavage assays using pSP1 and either *Sp*Cas9 or *St3*Cas9 with gRNAs as shown. M is the marker lane with bands (kbp): 10, 8.0, 6.0, 5.0, 4.0, 3.5, 3.0, 2.5, 2.0, 1.5, 1.0, 0.75, 0.5. Gels were run at different V/cm for different times, so the relative separation of the supercoiled (SC), nicked (OC) and linear (LIN) species differs. (**A**) DNA cleavage by *Sp*Cas9 using gRNA with unpaired 5′ modified gRNAs targeted to the 1147-1166 protospacer of pSP1 (Figure 3B). (**C**) DNA cleavage by *St3*Cas9 WT or nicking mutants using gRNA with unpaired 5′ modified gRNAs targeted to the 1025-1044 protospacer of pSP1 (Figure 4). (**C**) DNA cleavage by *Sp*Cas9 using RP-modified gRNAs targeted to the 1025-1044 protospacer of pSP1 (Figure 6A). N.B. The 5′ RP gels are presented in Figure 6B.

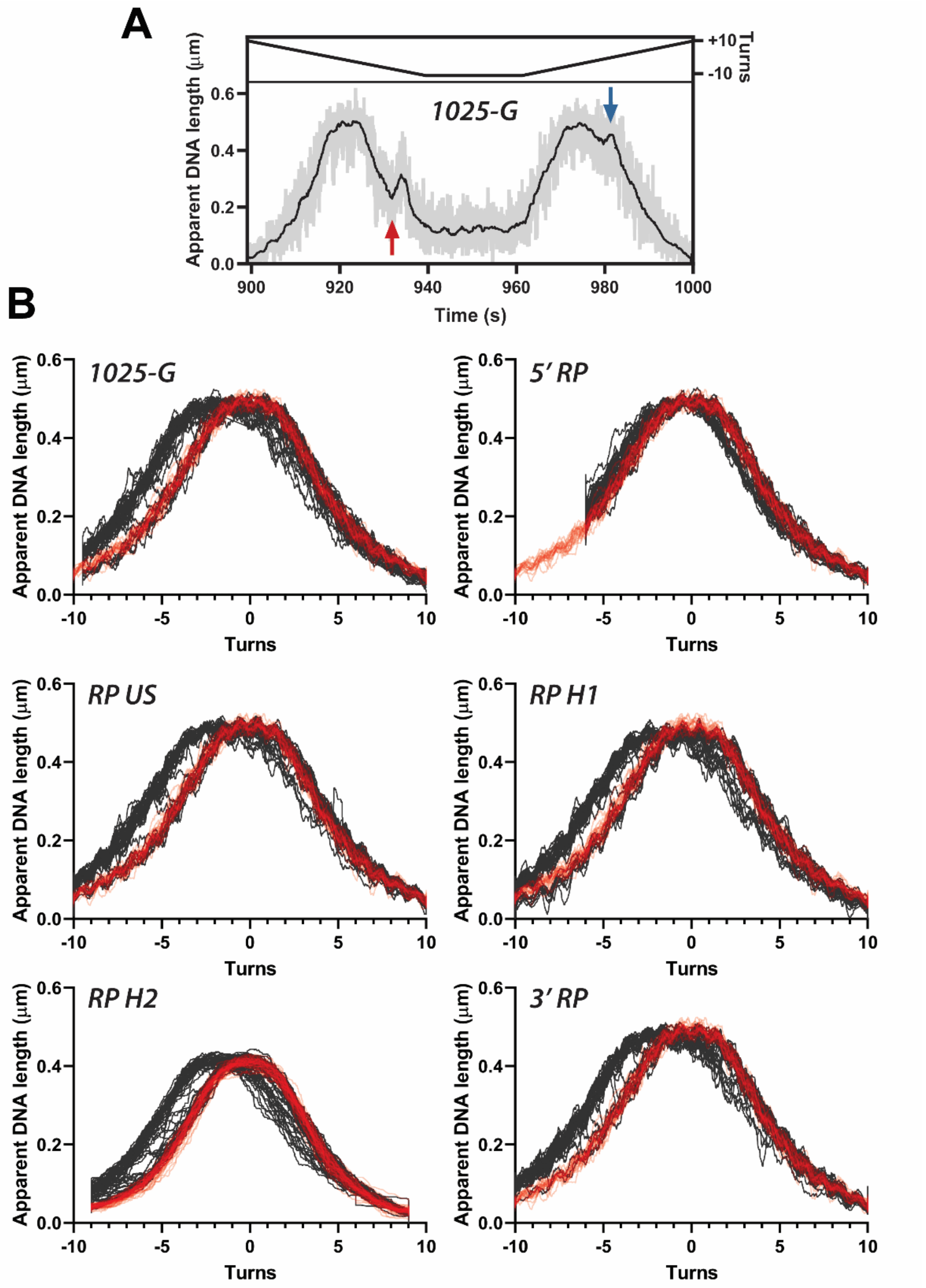


**Supplementary Figure 8.** R-loop cycling experiments in the MT assay. (**A**) Example of an R-loop cycling experiment at slow rotation speed (1 turn $s^{-1}$, 0.3 pN) in the presence of 1 nM *Sp*Cas9:1025-G gRNA. Raw DNA length recorded at 60 Hz (grey), data smoothed by a 2 Hz median filter (black). Events marked are R-loop formation at negative torque (red arrow) and dissociation at positive torque (blue arrow). The cycling between +10 and -10 turns is then repeated. (**B**) Overlay of repeated R-loop cycles (data smoothed by 2 Hz median filter) in the absence of Cas9 (red, *N* = 15), or in the presence of 1 nM *Sp*Cas9:gRNA as indicated (dark grey, *N* = 18-20). Rotation curve shifts (Figure 5E) were calculated by comparing the return parts of the curves where an R-loop is present from -6 to -4 turns. The RP H2 data was measured on a different DNA to the other gRNAs. The 5′ RP data was cycled between +10 and -6 to avoid surface interactions of the R-loop captured state.

| **Synthetic RNA** | **Sequence (5′ - 3′) – spacer sequence in lowercase.** | **Supplier** |
|---|---|---|
| *St3*Cas9 crRNA 1025 | cgcuaaagaggaagaggacaGUUUUAGAGCUGUGUUGUUUCG | Metabion |
| *Sp*Cas9 crRNA 1025 | cgcuaaagaggaagaggacaGUUUUAGAGCUAUGCUGUUUUG | Metabion |
| *Sp*Cas9 1166 | gcucacucauuaggcaccccGUUUUAGAGCUAGAAAUAGCAAGUUAAAAUAAGG CUAGUCCGUUAUCAACUUGAAAAAGUGGCACCGAGUCGGUGCUUUU | IDT Ultramer |
| *Sp*Cas9 1166-G | GgcucacucauuaggcaccccGUUUUAGAGCUAGAAAUAGCAAGUUAAAAUAAG GCUAGUCCGUUAUCAACUUGAAAAAGUGGCACCGAGUCGGUGCUUUU | IDT Ultramer |
| *Sp*Cas9 1166-GG | GGgcucacucauuaggcaccccGUUUUAGAGCUAGAAAUAGCAAGUUAAAAUAA GGCUAGUCCGUUAUCAACUUGAAAAAGUGGCACCGAGUCGGUGCUUUU | IDT Ultramer |
| *Sp*Cas9 1166-GGG | GGGgcucacucauuaggcaccccGUUUUAGAGCUAGAAAUAGCAAGUUAAAAUA AGGCUAGUCCGUUAUCAACUUGAAAAAGUGGCACCGAGUCGGUGCUUUU | IDT Ultramer |
| *Sp*Cas9 1166-UUU | UUUgcucacucauuaggcaccccGUUUUAGAGCUAGAAAUAGCAAGUUAAAAUA AGGCUAGUCCGUUAUCAACUUGAAAAAGUGGCACCGAGUCGGUGCUUUU | IDT ultramer |
| *St3*Cas9 1025-G | GcgcuaaagaggaagaggacaGUUUUAGAGCUGGGUACCCAGCGAGUUAAAAUA AGGCUUAGUCCGUACUCAACUUGAAAAGGUGGCACCGAUUCGGUGUUUU | IDT ultramer |
| *St3*Cas9 1025-GGG | GGGcgcuaaagaggaagaggacaGUUUUAGAGCUGGGUACCCAGCGAGUUAAAA UAAGGCUUAGUCCGUACUCAACUUGAAAAGGUGGCACCGAUUCGGUGUUUU | IDT ultramer |

**Supplementary table 1.** Synthetic crRNA and gRNA.

| **DNA substrates for IVT** | | | |
|---|---|---|---|
| **RNA** | **Plasmid** | **DNA sequence (5′ - 3′) – spacer in lowercase.** | **Primers pairs for PCR** |
| *St3*Cas9 tracrRNA | pCRISPR3 | CGAAACAACACAGCGAGTTAAAATAAGGCTTAGTCCGTACTCAACTTGAAAAGGTGGCACCGATTCGGTGTTTTT | TK-4 T7 Rev, TK-52 T7 |
| *Sp*Cas9 tracrRNA | pX330 | GTTTTAGAGCTAGAAATAGCAAGTTAAAATAAGGCTAGTCCGTTATCAACTTGAAAGTTTTAGAGCTAGAAATAGCAAGTAAGTGGCACCGAGTCG GTGCTTTTTT | TK-109 T7 Rev, TK-120 T7 |
| 1025-G | pD1301i-SP1 | cgctaaagaggaagaggacaGTTTTAGAGCTAGAAATAGCAAGTTAAAATAAGGCTAGTCCGTTATCAACTTGAAAAAGTGGCACCGAGTCGGTGCTTTTT T | oli#GM RP T7 (G)<br>oli#GM T7 Rev |
| 1025-GG | pD1301i-SP1 | As 1025-G | oli#GM RP T7 (GG)<br>oli#GM T7 Rev |
| 1025-GGG | pD1301i-SP1 | As 1025-G | oli#GM RP T7 (GGG)<br>oli#GM T7 Rev |
| 5′ RP | pEX-A2-g#GM SP1gRNA 5′ RP | TCTCCCTGAGCTTCAGGGAGcgctaaagaggaagaggacaGTTTTAGAGCTAGAAATAGCAAGTTAAAATAAGGCTAGTCCGTTATCAACTTGAAAAAGT GGCACCGAGTCGGTGCTTTTTT | oli#GM RP T7 (G)<br>oli#GM T7 Rev |
| 5′ RP $U_1$ | pEX 5′ RP U | TCTCCCTGAGCTTCAGGGAGTcgctaaagaggaagaggacaGTTTTAGAGCTAGAAATAGCAAGTTAAAATAAGGCTAGTCCGTTATCAACTTGAAAAAGT GGCACCGAGTCGGTGCTTTTTT | oli#GM RP T7 (G)<br>oli#GM T7 Rev |
| 5′ RP $U_2$ | pEX 5′ RP 2U | TCTCCCTGAGCTTCAGGGAGTTcgctaaagaggaagaggacaGTTTTAGAGCTAGAAATAGCAAGTTAAAATAAGGCTAGTCCGTTATCAACTTGAAAAAG TGGCACCGAGTCGGTGCTTTTTT | oli#GM RP T7 (G)<br>oli#GM T7 Rev |
| 5′ RP $U_4$ | pEX 5′ RP 4U | TCTCCCTGAGCTTCAGGGAGTTTTcgctaaagaggaagaggacaGTTTTAGAGCTAGAAATAGCAAGTTAAAATAAGGCTAGTCCGTTATCAACTTGAAAA AGTGGCACCGAGTCGGTGCTTTTTT | oli#GM RP T7 (G)<br>oli#GM T7 Rev |
| 5′ RP $U_8$ | pEX 5′ RP 8U | TCTCCCTGAGCTTCAGGGAGTTTTTTTTcgctaaagaggaagaggacaGTTTTAGAGCTAGAAATAGCAAGTTAAAATAAGGCTAGTCCGTTATCAACTTGA AAAAGTGGCACCGAGTCGGTGCTTTTTT | oli#GM RP T7 (G)<br>oli#GM T7 Rev |
| RP US | pEX-A2-g#GM SP1gRNA US RP | cgctaaagaggaagaggacaGTTTTAGAGCTCCCTGAGCTTCAGGGAGCAAGTTAAAATAAGGCTAGTCCGTTATCAACTTGAAAAAGTGGCACCGAGTC GGTGCTTTTTT | oli#GM SP1 T7 (G)<br>oli#GM T7 Rev |
| RP H1 | pEX-A2-g#GM SP1gRNA H RP_1 | cgctaaagaggaagaggacaGTTTTAGAGCTAGAAATAGCAAGTTAAAATAAGGCTAGTCCGTTATCAACTTGACTCCCTGAGCTTCAGGGAGAAAAGTG GCACCGAGTCGGTGCTTTTTT | oli#GM SP1 T7 (G)<br>oli#GM T7 Rev |
| RP H2 | pEX-A2-g#GM SP1gRNA H RP_2 | cgctaaagaggaagaggacaGTTTTAGAGCTAGAAATAGCAAGTTAAAATAAGGCTAGTCCGTTATCAACTCCCTGAGCTTCAGGGAGTGGCACCGAGTC GGTGCTTTTTT | oli#GM SP1 T7 (G)<br>oli#GM T7 Rev |
| RP 3′ | pEX-A2-g#GM SP1gRNA 3′ RP | cgctaaagaggaagaggacaGTTTTAGAGCTAGAAATAGCAAGTTAAAATAAGGCTAGTCCGTTATCAACTTGAAAAAGTGGCACCGAGTCGGTGCTCTC CCTGAGCTTCAGGGAGTTTTTT | oli#GM SP1 T7 (G)<br>oli#GM 3'RP T7 Rev |

| **Synthetic DNA primers** | | |
|---|---|---|
| | **DNA sequence (5′ - 3′) – T7 promoter in bold.** | Supplier |
| TK-4 T7 Rev | AAAAACACCGAATCGGTGCCAC | Metabion |
| TK-52 T7 | **TAATACGACTCACTATAGGG**CGAAACAACACAGCGAGTTAAAATAAGG | Metabion |
| TK-109 T7 Rev | AAAAAGCACCGACTCGGTGCCAC | Metabion |
| TK-120 T7 | **TAATACGACTCACTATAGGG**CAAAACAGCATAGCAAGTTAAAATAAGGCTAGTCCG | Metabion |
| oli#GM SP1 T7 (G) | GACA**CTAATACGACTCACTATAG**CGCTAAAGAGGAAGAGG | IDT |
| oli#GM SP1 T7 (GG) | GACA**CTAATACGACTCACTATAGG**CGCTAAAGAGGAAGAGG | IDT |
| oli#GM SP1 T7 (GG) | GACA**CTAATACGACTCACTATAGGG**CGCTAAAGAGGAAGAGG | IDT |
| oli#GM RP T7 (G) | GACA**CTAATACGACTCACTATAG**TCTCCCTGAGCTTCAGG | IDT |
| oli#GM T7 Rev | AAAAGCACCGACTCGGTGCC | IDT |
| oli#GM 3'RP T7 Rev | AAAACTCCCTGAAGCTCAGG | IDT |

**Supplementary Table 2.** DNA sequences of plasmids and synthetic DNA oligonucleotides used for PCR to make dsDNA substrates for *in vitro* transcription of gRNA and tracrRNA.

| *Sp*Cas9 | [Cas9] nM | Exponential fitted data IN events | | | | | | Exponential fitted data OUT events | | | | | |
|---|---|---|---|---|---|---|---|---|---|---|---|---|---|
| | | Torque pN nm | $\tau_{IN}$ (s) | | | | *N* | Torque pN nm | $\tau_{OUT}$ (s) | | | | *N* |
| **cr:tracr** | 20* | -6.0 | 100% | 21.02 ± 2.89 | | | 53 | +6.0 | 100% | 2.00 ± 0.27 | | | 53 |
| **cr:tracr** | 40* | -6.0 | 100% | 8.89 ± 1.34 | | | 44 | +6.0 | 100% | 2.25 ± 0.33 | | | 44 |
| **cr:tracr** | 80* | -6.0 | 100% | 4.15 ± 0.58 | | | 51 | +6.0 | 100% | 1.15 ± 0.16 | | | 51 |
| **cr:tracr** | 160* | -6.0 | 100% | 2.59 ± 0.34 | | | 59 | +6.0 | 100% | 1.23 ± 0.24 | | | 59 |
| **cr:tracr** | 20* | -5.0 | 100% | 18.73 ± 2.50 | | | 56 | +5.0 | 100% | 8.73 ± 1.16 | | | 56 |
| **cr:tracr** | 20* | -5.5 | 100% | 16.66 ± 2.33 | | | 51 | +5.5 | 100% | 4.15 ± 0.58 | | | 51 |
| **cr:tracr** | 20* | -6.0 | 100% | 13.41 ± 1.77 | | | 57 | +6.0 | 100% | 2.89 ± 0.38 | | | 57 |
| **cr:tracr** | 20* | -6.9 | 100% | 21.07 ± 2.86 | | | 54 | +6.9 | 100% | 0.71 ± 0.09 | | | 55 |
| **1G** | 10# | -6.2 | 100% | 3.02 ± 0.51 | | | 36 | +6.2 | 100% | 1.93 ± 0.32 | | | 36 |
| **2G** | 10# | -6.2 | 100% | 1.50 ± 0.23 | | | 43 | +6.2 | 100% | 2.85 ± 0.43 | | | 43 |
| **3G** | 10# | -6.2 | 100% | 2.21 ± 0.34 | | | 42 | +6.2 | 100% | 7.09 ± 1.09 | | | 42 |
| **5' RP** | 10# | -6.2 | 61% | 36.10 ± 8.28 | 39% | 3.57 ± 0.82 | 19 | +6.2 | 36% | 0.07 ± 0.02 s | 64% | 8.72 ± 2.00 | 19 |
| **US RP** | 10# | -6.2 | 100% | 18.68 ± 3.74 | | | 25 | +6.2 | 100% | 0.40 ± 0.08 s | | | 25 |
| **H1 RP** | 10# | -6.2 | 100% | 10.90 ± 2.73 | | | 16 | +6.2 | 100% | 0.64 ± 0.16 s | | | 16 |
| **H2 RP** | 10# | -6.2 | 100% | 5.28 ± 1.04 | | | 26 | +6.2 | 54% | 0.39 ± 0.08 s | 46% | 5.91 ± 1.16 | 26 |
| **3' RP** | 10# | -6.2 | 100% | 16.81 ± 3.86 | | | 19 | +6.2 | 100% | 0.61 ± 0.14s | | | 19 |

| *St3*Cas9 | [Cas9] nM | Exponential fitted data IN events | | | | | | Exponential fitted data OUT events | | | | | |
|---|---|---|---|---|---|---|---|---|---|---|---|---|---|
| | | Torque pN nm | $\tau_{IN}$ (s) | | | | *N* | Torque pN nm | $\tau_{OUT}$ (s) | | | | *N* |
| **cr:tracr** | 1 | -6.0 | 100% | 8.03 ± 1.07 | | | 56 | +6.0 | 100% | 1.16 ± 0.15 | | | 55 |
| **cr:tracr** | 2 | -6.0 | 100% | 5.82 ± 0.80 | | | 53 | +6.0 | 100% | 0.93 ± 0.12 | | | 53 |
| **cr:tracr** | 4 | -6.0 | 100% | 1.28 ± 0.17 | | | 58 | +6.0 | 100% | 1.19 ± 0.15 | | | 58 |
| **cr:tracr** | 8 | -6.0 | 100% | 0.68 ± 0.08 | | | 64 | +6.0 | 100% | 1.58 ± 0.20 | | | 63 |
| **cr:tracr** | 1 | -5.0 | 100% | 9.76 ± 1.53 | | | 41 | +5.0 | 100% | 5.33 ± 0.83 | | | 40 |
| **cr:tracr** | 1 | -5.5 | 100% | 7.41 ± 1.03 | | | 51 | +5.5 | 100% | 1.83 ± 0.25 | | | 50 |
| **cr:tracr** | 1 | -6.1 | 100% | 7.33 ± 0.94 | | | 60 | +6.1 | 100% | 2.02 ± 0.26 | | | 58 |
| **cr:tracr** | 1 | -7.0 | 100% | 6.02 ± 0.79 | | | 58 | +7.0 | 100% | 0.68 ± 0.09 | | | 58 |

| *St3*Cas9 D31A | [Cas9] nM | Exponential fitted data IN events | | | | | | Exponential fitted data OUT events | | | | | |
|---|---|---|---|---|---|---|---|---|---|---|---|---|---|
| | | Torque pN nm | $\tau_{IN}$ (s) | | | | *N* | Torque pN nm | $\tau_{OUT}$ (s) | | | | *N* |
| **cr:tracr** | 1 | -6.0 | 100% | 7.89 ± 1.11 | | | 50 | +6.0 | 100% | 1.94 ± 0.27 | | | 50 |
| **cr:tracr** | 2 | -6.0 | 100% | 3.31 ± 0.45 | | | 53 | +6.0 | 100% | 1.94 ± 0.26 | | | 51 |
| **cr:tracr** | 4 | -6.0 | 100% | 2.07 ± 0.28 | | | 57 | +6.0 | 100% | 1.87 ± 0.25 | | | 57 |
| **cr:tracr** | 8 | -6.0 | 100% | 1.01 ± 0.13 | | | 58 | +6.0 | 100% | 2.48 ± 0.33 | | | 57 |
| **cr:tracr** | 1 | -5.0 | 100% | 5.29 ± 0.78 | | | 46 | +5.0 | 100% | 3.47 ± 0.51 | | | 46 |
| **cr:tracr** | 1 | -5.5 | 100% | 10.45 ± 1.50 | | | 49 | +5.5 | 100% | 3.03 ± 0.42 | | | 49 |
| **cr:tracr** | 1 | -6.1 | 75% | 11.49 ± 1.58 | 25% | 1.24 ± 0.17 | 53 | +6.1 | 100% | 2.93 ± 0.40 | | | 53 |
| **cr:tracr** | 1 | -7.0 | 100% | 8.90 ± 1.17 | | | 57 | +7.0 | 100% | 2.10 ± 0.28 | | | 57 |

| *St3*Cas9 N891A | [Cas9] nM | Exponential fitted data IN events | | | | | | Exponential fitted data OUT events | | | | | |
|---|---|---|---|---|---|---|---|---|---|---|---|---|---|
| | | Torque pN nm | $\tau_{IN}$ (s) | | | | *N* | Torque pN nm | $\tau_{OUT}$ (s) | | | | *N* |
| **cr:tracr** | 1 | -6.0 | 100% | 7.48 ± 1.14 | | | 43 | +6.0 | 100% | 0.78 ± 0.11 | | | 43 |
| **cr:tracr** | 2 | -6.0 | 100% | 2.54 ± 0.43 | | | 34 | +6.0 | 100% | 2.34 ± 0.39 | | | 31 |
| **cr:tracr** | 4 | -6.0 | 100% | 2.10 ± 0.35 | | | 36 | +6.0 | 100% | 2.22 ± 0.37 | | | 36 |
| **cr:tracr** | 8 | -6.0 | 100% | 1.45 ± 0.20 | | | 51 | +6.0 | 100% | 2.08 ± 0.29 | | | 50 |
| **cr:tracr** | 1 | -4.7 | 100% | 5.29 ± 0.71 | | | 56 | +4.7 | 100% | 4.56 ± 0.61 | | | 53 |
| **cr:tracr** | 1 | -5.3 | 100% | 4.67 ± 0.66 | | | 51 | +5.3 | 100% | 1.34 ± 0.20 | | | 48 |
| **cr:tracr** | 1 | -5.8 | 100% | 5.53 ± 0.74 | | | 55 | +5.8 | 100% | 1.45 ± 0.20 | | | 55 |
| **cr:tracr** | 1 | -6.5 | 79% | 5.24 ± 0.68 | 21% | 0.31 ± 0.04 | 59 | +6.5 | 100% | 0.85 ± 0.11 | | | 55 |

| *St3*dCas9 | [Cas9] nM | Exponential fitted data IN events | | | | | | Exponential fitted data OUT events | | | | | |
|---|---|---|---|---|---|---|---|---|---|---|---|---|---|
| | | Torque pN nm | $\tau_{IN}$ (s) | | | | *N* | Torque pN nm | $\tau_{OUT}$ (s) | | | | *N* |
| **cr:tracr** | 1 | -6.0 | 100% | 7.79 ± 1.00 | | | 61 | +6.0 | 100% | 2.85 ± 0.37 | | | 60 |
| **cr:tracr** | 2 | -6.0 | 100% | 3.71 ± 0.52 | | | 50 | +6.0 | 100% | 1.62 ± 0.23 | | | 50 |
| **cr:tracr** | 4 | -6.0 | 64% | 3.45 ± 0.49 | 36% | 0.24 ± 0.03 | 48 | +6.0 | 100% | 2.49 ± 0.36 | | | 48 |
| **cr:tracr** | 8 | +6.0 | 100% | 1.70 ± 0.23 | | | 57 | +6.0 | 100% | 2.17 ± 0.29 | | | 56 |
| **cr:tracr** | 1 | +4.7 | 100% | 8.87 ± 1.32 | | | 45 | +4.7 | 100% | 10.07 ± 1.49 | | | 44 |
| **cr:tracr** | 1 | +5.2 | 100% | 8.31 ± 1.19 | | | 49 | +5.2 | 100% | 3.76 ± 0.54 | | | 49 |
| **cr:tracr** | 1 | +5.7 | 100% | 7.69 ± 1.10 | | | 49 | +5.7 | 100% | 21.97 ± 0.28 | | | 49 |
| **cr:tracr** | 1 | +6.5 | 100% | 7.89 ± 1.11 | | | 51 | +6.5 | 100% | 1.84 ± 0.25 | | | 50 |

**Supplementary Table 3.** Time constants from the fits to the IN or OUT times measured from the magnetic tweezers assay (Figures 3D, 5G, 5H, and Supplementary Figure 3). Data plotted in Figures 2D, 2E, 3E, and 5I. 100% indicates single exponential, two percentages indicate the amplitudes of the fast and slow phases of a double exponential. *SpCas9 preparation 2014-09-23. #New England Biolabs *Sp*Cas9 M0386. Errors are standard errors based on *N* values.

| (± SD) | $k_{R+loop}$ ($s^{+1}$) | $k_a$ ($s^{-1}$) | $k_b$ ($s^{-1}$) | $k_{ini}$ ($s^{-1}$) | $k_{ini2}$ ($s^{-1}$) | *N* |
|---|---|---|---|---|---|---|
| **Figure 3H** | | | | | | |
| **1025+G** | N/A | 0.0484 ± 0.0068 | 0.10545 ± 0.00501 | 0.0211 ± 0.0107 | 0.0091 ± 0.0013 | 4 |
| **1025+GG** | N/A | 0.0214 ± 0.0006 | 0.00392 ± 0.00041 | 0.0067 ± 0.0059 | 0.0007 ± 0.0002 | 4 |
| **1025+GGG** | N/A | 0.0216 ± 0.0014 | 0.00098 ± 0.00019 | 0.0115 ± 0.0033 | 0.0005 ± 0.0002 | 3 |
| **Figure 6C** | | | | | | |
| **1025+G** | N/A | 0.0987 ± 0.0756 | 0.1041 ± 0.0132 | 0.0047 ± 0.0017 | 0.0062 ± 0.0008 | 3 |
| **RP US** | N/A | 0.0883 ±0.0085 | 0.1281 ± 0.0180 | 0.0067 ± 0.0012 | 0.0100 ± 0.0037 | 3 |
| **RP H1** | N/A | 0.1156 ± 0.0177 | 0.1565 ± 0.0314 | 0.0057 ± 0.0027 | 0.0107 ± 0.0039 | 3 |
| **RP H2** | N/A | 0.0871 ± 0.0075 | 0.1603 ± 0.0271 | 0.0067 ± 0.0020 | 0.0098 ± 0.0032 | 3 |
| **3’ RP** | N/A | 0.0990 ± 0.0056 | 0.1239 ± 0.0107 | 0.0045 ± 0.0022 | 0.0078 ± 0.0019 | 3 |
| **Figure 4F** | | | | | | |
| **1166** | N/A | 0.071 ± 0.031 | 0.722 ± 0.430 | 0.007 ± 0.004 | 0.106 ± .068 | 3 |
| **1166+G** | N/A | n.d. | n.d. | n.d. | n.d. | 3 |
| **1166+GG** | N/A | 0.089 ± 0.041 | 0.042 ± 0.015 | 0.018 ± 0.011 | 0.009 ± 0.001 | 3 |
| **1166+GGG** | N/A | 0.009 ± 0.003 | 0.021 ± 0.003 | 0.007 ± 0.001 | 0.009 ± 0.004 | 3 |
| **1166+GGG (2)** | 0.017 ± 0.004 | 0.14 fixed | 0.048 ± 0015 | 0.114 ± 0.05 | 0.015 ± 0.007 | 3 |
| **1166+UUU** | N/A | 0.129 ± 0.037 | 0.019 ± 0.002 | 0.019 ± .007 | 0.005 ± 0.002 | 3 |

**Supplementary Table 4.** Apparent rate constants from the numerical integration fits of the DNA cleavage data in Figures 3F, 4B, 4E and 6A to the models in Supplementary Figure 6. Each repeat was fitted separately, and the fitted parameters averaged. Errors are standard deviation.
